# FUS modulates the level of ribosomal RNA modifications by regulating a subset of snoRNA expression

**DOI:** 10.1101/2022.11.09.515592

**Authors:** Kishor Gawade, Patrycja Plewka, Sophia J Häfner, Anders H Lund, Virginie Marchand, Yuri Motorin, Michal W Szczesniak, Katarzyna D Raczynska

## Abstract

FUS is a multifunctional protein involved in many aspects of RNA metabolism, including transcription, splicing, translation, miRNA processing, and replication-dependent histone gene expression. In this paper, we show that FUS depletion results in differential expression of numerous small nucleolar RNAs (snoRNAs) that guide 2’-O methylation (2’-O-Me) and pseudouridylation of specific positions in ribosomal RNAs (rRNAs) and small nuclear RNAs (snRNAs). Using RiboMeth-seq and HydraPsiSeq for the profiling of 2’-O-Me and pseudouridylation status of rRNA species, we demonstrated considerable hypermodification at several sites in HEK293T and SH-SY5Y cells with FUS knockout (FUS KO) compared to wild-type cells. We observed a similar direction of changes in rRNA modification in differentiated SH-SY5Y cells with the FUS mutation (R495X) related to the severe disease phenotype of amyotrophic lateral sclerosis (ALS). Furthermore, the pattern of modification of some rRNA positions was correlated with the abundance of corresponding guide snoRNAs in FUS KO and FUS R495X cells. Our findings reveal a new role for FUS in modulating the modification pattern of rRNA molecules, that in turn might generate ribosome heterogeneity and constitute a fine-tuning mechanism for translation efficiency/fidelity. Therefore, we suggest that increased levels of 2’-O-Me and pseudouridylation at particular positions in rRNAs from cells with the ALS-linked FUS mutation may represent a possible new translation-related mechanism that underlies disease development and/or progression.

## INTRODUCTION

FUS belongs to the FET family of proteins that includes three highly conserved, abundant, and ubiquitously expressed RNA-binding proteins: FUS, EWSR1, and TAF15 (1). FUS is a nucleocytoplasmic shuttling protein predominantly localized to the nucleus, but it can also be found in the cytoplasm and was shown to participate in mRNA transport and translation (2,3). It also binds to both ssDNA (single-stranded DNA) and dsDNA (double-stranded DNA), facilitating DNA annealing and D-loop formation and thus mediating genomic maintenance, DNA recombination, and DNA repair (4–7). FUS regulates critical steps in RNA metabolism, including transcription, splicing, and alternative splicing (reviewed in (8)). Furthermore, FUS is involved in replication-dependent histone gene expression, microRNA biogenesis, and function (9–11). In particular, numerous FUS mutations have been found in familial forms of amyotrophic lateral sclerosis (ALS) and frontotemporal lobar degeneration (FTLD), indicating an essential role for this protein in neurodegenerative diseases (1,8,12– 16).

Small nucleolar ribonucleoproteins (snoRNPs) are nucleolus-localized complexes comprising small nucleolar RNAs (snoRNAs) associated with highly conserved core proteins. snoRNPs are categorized into two major classes, referred to as “box C/D” and “box H/ACA” snoRNPs, depending on the presence of “C/D” or “H/ACA” sequence motifs in the snoRNAs. Most snoRNPs function in precursor rRNA (pre-rRNA) maturation by introducing sequence-specific modifications guided by snoRNAs. C/D snoRNPs are responsible for site-specific 2’*-*O*-*ribose methylation (2’-O-Me), while H/ACA snoRNPs catalyze the conversion of uridine to pseudouridine (ψ). Additionally, members of another subgroup of snoRNPs – Cajal body-specific snoRNPs (scaRNPs) – modify the RNA component of U snRNPs (small nuclear RNAs), a class of non-coding RNAs involved mainly in the splicing of pre-mRNAs. Two enzymes are responsible for the catalytic activity of snoRNPs: the 2’-O-methyltransferase fibrillarin (FBL) and the pseudouridine synthase dyskerin (DKC1) (17). Interestingly, some snoRNAs are pairing with pre-rRNAs but are not involved in any chemical modifications. Instead, they play a crucial role in pre-rRNA processing, such as U3 and U8 snoRNAs (17,18). Moreover, many “orphan” snoRNAs have been identified in higher eukaryotes. These snoRNAs possess either the C/D box or H/ACA box motif but do not correspond to any target RNAs, suggesting that they may play different functions in cells (19).

2’-O-Me and pseudouridylation are the most abundant chemical modifications identified in ribosomal RNAs (20). The modified sites are within the functionally important domains and also on the ribosome periphery and may influence the proper folding and translation efficiency of ribosomes as reported by the previous studies (21–25). Recently, Jansson et al. reported that differential 2’-O-Me is a source of ribosomal heterogeneity and that Myc-induced changes in one methylation site, 18S-Cm174, can result in altered expression of specific mRNAs, which correlates with cell phenotype changes (25). A study by Krogh et al. suggests that the methylation status of 18S-Cm1440 differentiates ribosomes and influences the affinity of translation factors involved in the recruitment of mRNAs (24). There is also evidence to confirm that pseudouridine affects ribosomal function *in vivo*, and defects in these functions may lead to impaired ribosome-ligand interactions, as observed in X-linked dyskeratosis congenita (X-DC) (22). To summarize, earlier reports highlight that methylation and pseudouridylation of ribosomal RNAs are key players in producing specialized ribosomes that could have tissue, organ, or disease etiology-specific expression. Moreover, previous studies identified that 2’-O-Me and pseudouridine modifications are also necessary for proper secondary structure and binding of snRNP components (26). An earlier study by Yu et al. confirmed that modified nucleotides within the 5’ end of U2 snRNA are necessary to form functional splicing complexes (27). Furthermore, as reported by Nagasawa et al., pseudouridine modification at position U89 of U2 snRNA depends on scaRNA1 levels and affects the splicing of mRNAs involved in embryonic development (28).

In our study, small RNA sequencing revealed that many C/D box snoRNAs, H/ACA box snoRNAs, and scaRNAs are differentially expressed in FUS knockout (FUS KO) cells. This prompted us to look for possible changes in 2’-O-Me and pseudouridylation status of rRNAs and snRNAs, as these modifications are mostly catalyzed by snoRNP complexes. We used RiboMeth-seq (RMS) and HydraPsiSeq, which are 2’-O-Me and pseudouridine detection and quantification methods, in which alkaline hydrolysis and hydrazine*/*aniline cleavage were coupled to high throughput sequencing, respectively (29,30). We studied HEK293T FUS KO cells and SH-SY5Y (neuroblastoma) FUS KO cells as well as SH-SY5Y cells with FUS R495X mutation associated with the severe disease phenotype of ALS. Our data reveal positions in 18S and 28S rRNAs, which are methylated and pseudouridylated on a significantly higher proportion of the ribosomes in FUS KO and FUS R495X cells. These positions are located in the functional domains and periphery of the small subunit (SSU: 18S rRNA) and large subunit (LSU: 5S, 5.8S, and 28S rRNA). These sites are solvent accessible and demonstrate ribosome heterogeneity at the level of rRNA modifications in different cell lines and other physiological conditions, like proliferation and differentiation. Our report suggests a novel role for FUS in modulating the status of rRNA modifications by regulating the expression of the subset of snoRNAs. It also proposes that ALS-associated FUS mutations might lead to increased 2’-O-Me and pseudouridylation of 18S and 28S rRNAs at specific sites, which may result in decreased translational fidelity or reduced translation of a specific set of mRNAs as observed in latest studies with particular snoRNA knockout or decrease in the steady state level of snoRNAs by introducing mutations in the biogenesis machinery (25,31,32). Moreover, as multiple scaRNAs have been found to be differentially expressed in FUS KO cells, and some pseudouridylation sites in snRNAs were altered in these cells and in FUS R495X mutant, this could partially explain RNA splicing defects reported in FUS depleted cells and caused by ALS-associated FUS mutations (33–35).

## RESULTS

### FUS regulates the level of snoRNAs in human cells

FUS is known to regulate microRNA biogenesis and microRNA-mediated gene silencing (10,11). It also interacts with spliceosomal U snRNAs (35–38) and we previously reported the interaction of FUS with U7 snRNA (9,16). Moreover, many microRNAs and other small RNAs, including snoRNAs, were differentially expressed in sporadic ALS patients compared to healthy age-matched controls (39). Based on these, we wanted to explore whether FUS is involved in the regulation of expression of snoRNAs. For this purpose, we performed preliminary experiment of high-throughput sequencing of small RNAs (small RNA-seq) isolated from SH-SY5Y wild-type (WT) and FUS KO cells, both proliferating and differentiated to neuron-like cells. PCA (principal component analysis) was performed to identify whether there was difference in sample clustering, as observed in Supplementary Fig. S1A, B; WT and FUS KO samples formed separate clusters. Next, we counted the reads assigned according to GTF.attrType = “gene_biotype”, in FUS KO differentiated cells higher proportion of reads were assigned to snoRNAs as compared to wild-type cells (Supplementary Fig. S1C, D). As shown in Fig. 1, Supplementary Table S1, and Supplementary Fig. S2, we identified C/D box snoRNAs, H/ACA box snoRNAs, and scaRNAs that were differentially expressed between FUS KO and WT cells, both in proliferating and differentiated cells. They include both unique snoRNAs as well as some snoRNAs from genetically-imprinted C/D box snoRNA families, like SNORD113, SNORD114, and SNORD116 (Supplementary Table S1), suggesting that FUS may directly bind to individual snoRNAs or may be involved in the regulation of these snoRNAs indirectly, by controlling the expression of their host genes.

**Fig. 1.**
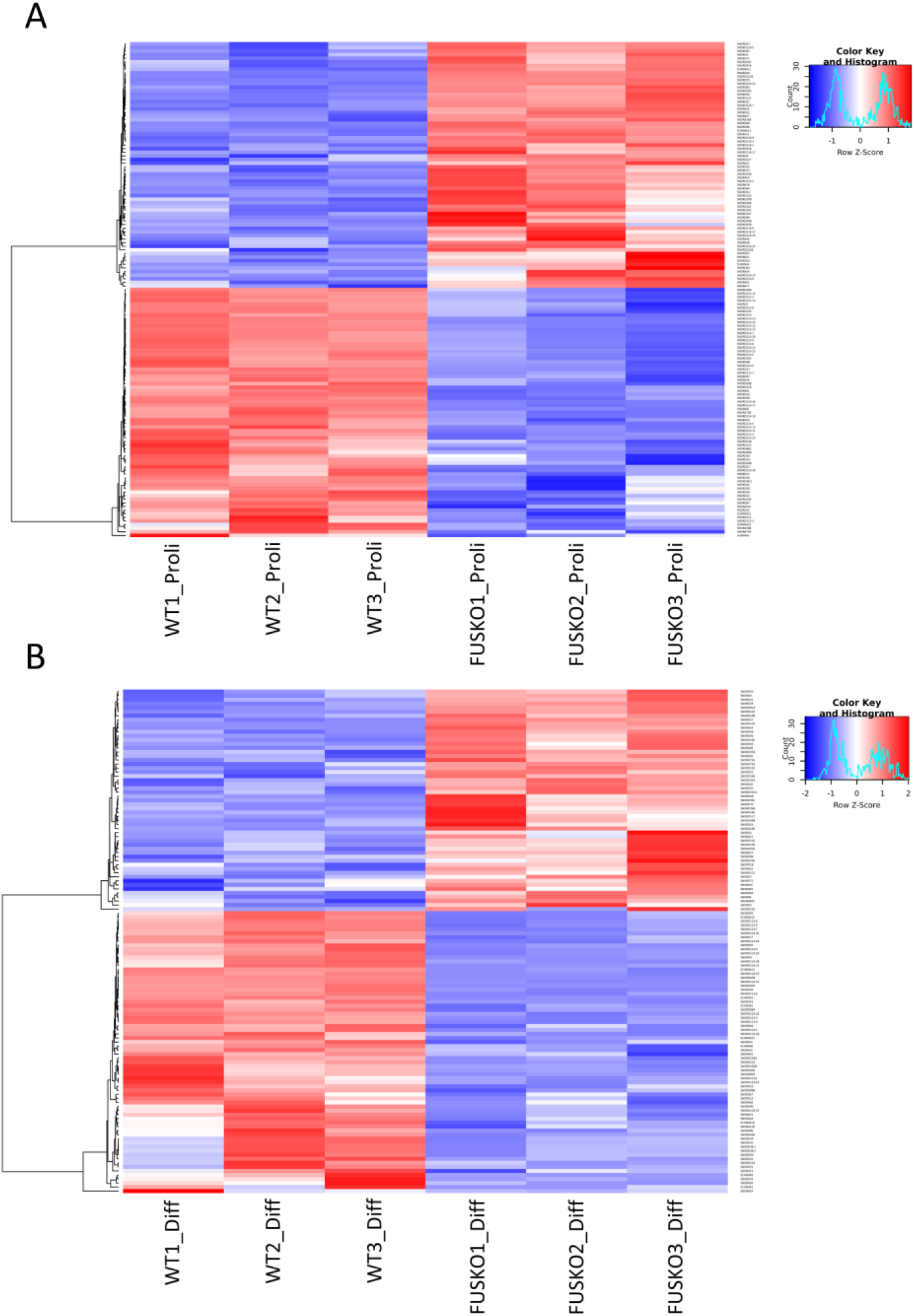
Differentially expressed snoRNAs in the proliferating (Proli) (A) and differentiated (Diff) (B) SH-SY5Y WT and FUS KO cells identified by small RNA sequencing. DESeq normalized read count was used to generate a heatmap for differentially expressed genes with Heatmap.2 function of gplots R package. Padj (p-adjusted value cut off <0.05). The blue and red colors show a low and high level of expression, respectively.

### FUS depletion and FUS R495X mutation affect 2’-O-Me levels at particular positions in 18S and 28S rRNAs

The results obtained prompted us to inquire the main question of this project, whether the changes in guide RNA levels will eventually modulate the 2’-O-Me and pseudouridylation levels of SSU and LSU ribosomal RNA subunits. To gain insight into the impact of FUS depletion on global 2’-O-Me signatures, we decided to analyze WT and FUS KO cells from two different cell lines: HEK293T and SH-SY5Y proliferating and differentiated. Furthermore, we analyzed SH-SY5Y cells with the ALS-linked FUS R495X mutation differentiated with retinoic acid for 10 days. Using RiboMeth-seq, we determined the methylation pattern of previously reported 68 positions in 28S rRNA, 41 sites in 18S rRNA, and two positions in 5.8S rRNA in all cell lines analyzed (Supplementary Table S2). A methylation score (MethScore) was calculated for each position to assess the fraction of rRNA molecules modified at a given position in all FUS KO cells, FUS R495X cells and WT cells. The vast majority of sites were unaffected by the FUS knockout or the FUS R495X mutation in both rRNA subunits and represented almost saturated 2’-O-Me in all cell lines analyzed (Supplementary Table S2, Supplementary Fig. S3A). The affected sites (MethScore difference > 0.05) corresponded mainly to higher 2’-O-Me levels in FUS KO and FUS R495X cells in comparison to WT cells (Table 1A-C, Fig. 2A). The lower proportion of 2’-O-methylation was observed only in the case of 18S-Am576 in FUS KO SH-SY5Y proliferating cells and 28S-Am1323 in differentiated SH-SY5Y FUS KO and FUS R495X cells. In 18S rRNA, the methylation status of two positions, 18S-Um354 and 18S-Cm1272, was consistently and noticeably higher in all FUS KO cells and FUS R495X cells compared to WT. Moreover, the 18S-Gm436 position was sensitive to FUS depletion and mutation in neuroblastoma cells only. Comparison of 28S rRNA methylation profiles revealed that two positions, 28S-Am2401 and 28S-Gm2876, were concordantly hypermethylated in HEK293T FUS KO cells and differentiated SH-SY5Y FUS KO and FUS R495X cells (Table 1A-C, Fig. 2A). Interestingly, 28S rRNA methylation profile was unchanged in SH-SY5Y proliferating cells (Table 1B). These results support the notion that FUS disturbances (FUS depletion or ALS-linked mutations) can affect the methylation status of specific rRNA positions, with two sites: 18S-Um354 and 18S-Cm1272, appearing to be commonly responsive to FUS mutations. Furthermore, most FUS sensitive 2’-O-Me sites exhibited methylation scores lower than 0.65 (Table 1A-C), suggesting that mainly sub-stoichiometric modifications are influenced explicitly by FUS mutations (29).

**Table 1A.**
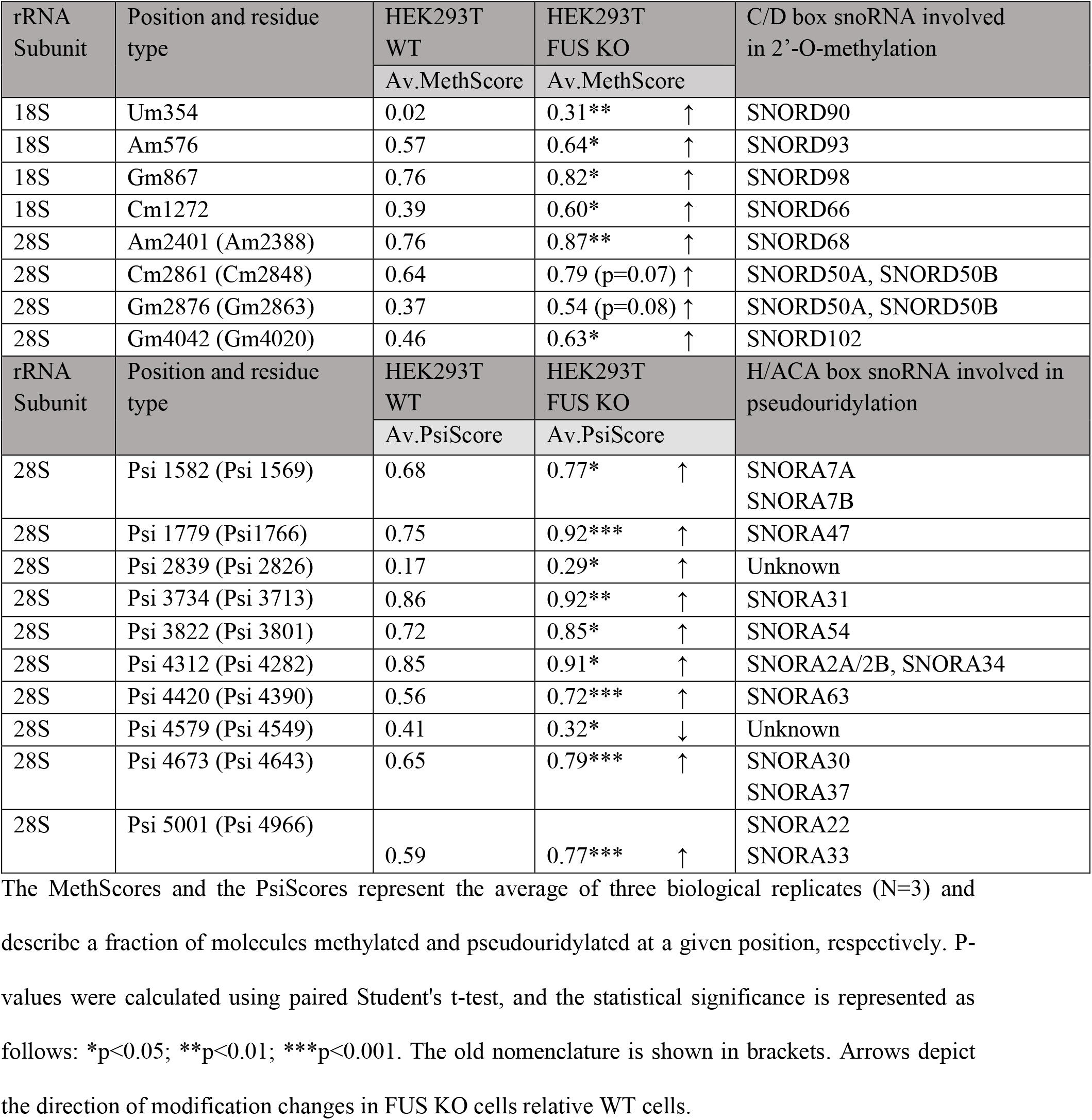
2’-O-Me and pseudouridylation positions changed in 18S and 28S ribosomal RNAs in HEK293T WT and FUS KO cell lines.

**Table 1B.** 2’-O-methylation and pseudouridylation positions changed in 18S and 28S ribosomal RNAs in SH-SY5Y WT and FUS KO cell lines in proliferating (Proli) conditions.

| rRNA Subunit | Position and residue type | SH-SY5Y WT Proli | SH-SY5Y FUS KO Proli | C/D box snoRNA involved in 2'-O-methylation |
| --- | --- | --- | --- | --- |
|  |  | Av.MethScore | Av.MethScore |  |
| 18S | Um354 | 0.44 | 0.61 (p=0.08) ↑ | SNORD90 |
| 18S | Gm436 | 0.47 | 0.62** ↑ | SNORD100 |
| 18S | Am576 | 0.19 | 0.09* ↓ | SNORD93 |
| 18S | Cm1272 | 0.58 | 0.71* ↑ | SNORD66 |
| rRNA Subunit | Position and residue type | SH-SY5Y WT Proli | SH-SY5Y FUS KO Proli | H/ACA box snoRNA involved in pseudouridylation |
|  |  | Av.PsiScore | Av.PsiScore |  |
| 18S | Psi 681 | 0.82 | 0.73* ↓ | Unknown |
| 28S | Psi 5001 (Psi 4966) | 0.69 | 0.78* ↑ | SNORA22<br>SNORA33 |

The MethScores and the PsiScores represent the average of three biological replicates (N=3) and describe a fraction of molecules methylated and pseudouridylated at a given position, respectively. P-values were calculated using paired Student's t-test, and the statistical significance is represented as follows: \*p<0.05; \*\*p<0.01. The old nomenclature is shown in brackets. Arrows depict the direction of modification changes in FUS KO cells relative WT cells.

**Table 1C.** 2’-O-methylation and pseudouridylation positions changed in 18S and 28S ribosomal RNAs in SH-SY5Y WT, FUS KO and FUS R495X cells differentiated (Diff) for 10 days using RA (retinoic acid).

| rRNA Subunit | Position and residue type | SH-SY5Y WT Diff | SH-SY5Y FUS KO Diff | SH-SY5Y FUS R495X Diff | C/D box snoRNA involved in 2'-O-methylation |
| --- | --- | --- | --- | --- | --- |
|  |  | Av.MethScore | Av.MethScore | Av.MethScore |  |
| 18S | Um354 | 0.59 | 0.71* ↑ | 0.85** ↑ | SNORD90 |
| 18S | Gm436 | 0.61 | 0.74* ↑ | 0.86** ↑ | SNORD100 |
| 18S | Cm1272 | 0.69 | 0.81* ↑ | 0.81* ↑ | SNORD66 |
| 18S | Cm1440 | 0.41 | 0.52 (p=0.14) ↑ | 0.56 | SNORD125 |
|  |  |  |  | (0.066) ↑ |  |
| **28S** | Am1323 (Am1310) | 0.63 | 0.45<br>(p=0.1) ↓ | 0.41<br>(p=0.055) ↓ | SNORD126 |
| 28S | Cm1881 (Cm1868) | 0.68 | 0.77 (p=0.15) ↑ | 0.81<br>(p=0.08) ↑ | SNORD48 |
| 28S | Am2401 (Am2388) | 0.90 | 0.94* ↑ | 0.96* ↑ | SNORD68 |
| 28S | Am2787 (Am2774) | 0.86 | 0.89 (p=0.07) | 0.92** ↑ | SNORD99 |
| 28S | Gm2876 (Gm2863) | 0.83 | 0.91** ↑ | 0.94** ↑ | SNORD50A,<br>SNORD50B |
| 28S | Am3867 (Am3846) | 0.80 | 0.87* ↑ | 0.90* ↑ | SNORD92 |
| 28S | Cm4456 (Cm4426) | 0.91 | 0.97* ↑ | 0.98* ↑ | SNORD49A,<br>SNORD49B |
| rRNA<br>Subunit | Position and residue<br>type | SH-SY5Y WT<br>Diff | SH-SY5Y FUS<br>KO Diff | SH-SY5Y<br>FUS R495X<br>Diff | H/ACA box<br>snoRNA involved in<br>pseudouridylation |
|  |  | Av.PsiScore | Av.PsiScore | Av.PsiScore |  |
| 18S | Psi 897 | -0.08 | -0.0066* ↓ | 0.14*** ↑ | SNORA44 |
| 28S | Psi 2839 (Psi 2826) | 0.50 | 0.57* ↑ | 0.61*** ↑ | Unknown |
| 28S | Psi 2843 (Psi 2830) | -3.9 | -3.6 (p=0.15) | -2.9** ↑ | Unknown |
| 28S | Psi 4579 (Psi 4549) | 0.78 | 0.79 (p=0.29) | 0.87*** ↑ | Unknown |
| 28S | Psi 4636 (Psi 4606) | 0.51 | 0.42** ↓ | 0.43** ↓ | SNORA81 |
| 28S | Psi 4973 (Psi 4938) | -1.03 | -0.95 (p=0.63) | -0.78* ↑ | SNORA43 |

The MethScores and the PsiScores represent the average of three biological replicates (N=3) and describe a fraction of molecules methylated and pseudouridylated at a given position, respectively. P-values were calculated using paired Student's t-test, and the statistical significance is represented as follows: \*p<0.05; \*\*p<0.01; \*\*\*p<0.001. The old nomenclature is shown in brackets. Arrows depict the direction of modification changes in FUS KO and FUS R495X cells relative WT cells. Negative values mean that these residues are not protected against hydrazine/aniline treatment, hence they are not pseudouridylated.

**Fig. 2.**
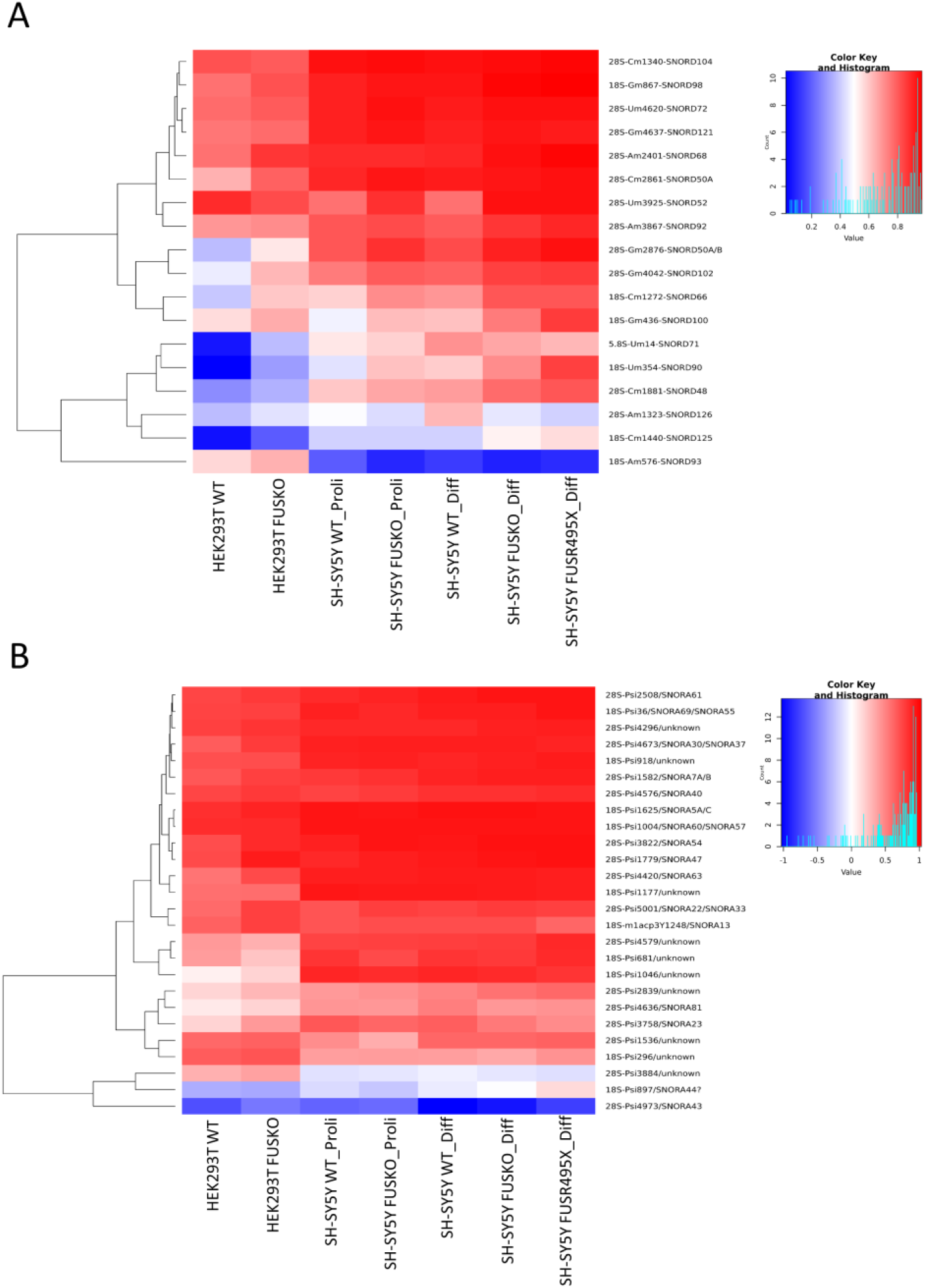
Differential heatmaps for variations in 2’-O-Me (A) and pseudouridylation (B) levels in all cells analyzed (HEK293T WT and FUS KO, SH-SY5Y proliferating (Proli) WT and FUS KO, SH-SY5Y differentiated (Diff) WT, FUS KO and FUS R495X), representing the average of three biological replicates (N=3) from each cell line. Heatmaps represent sites with at least a 10% (0.1) difference in 2’-O-Me or pseudouridylation between cell-types. The blue and red colors show hypomodification and hypermodification, respectively.

In the next step, we checked the expression levels of selected box C/D snoRNAs that guide the 2’-O-Me rRNA modification of sites affected in FUS KO or FUS R495X cells. For this purpose, we used previously described small RNA-seq data from neuroblastoma WT and FUS KO cells (proliferating and differentiated) and supplemented them by RT-qPCR approach done in all cells analyzed (HEK293T WT and FUS KO, SH-SY5Y proliferating (Proli) WT and FUS KO, SH-SY5Y differentiated (Diff) WT, FUS KO and FUS R495X) (Supplementary Table S1, Fig. 3A, C, Fig. 4). In HEK293T FUS KO cells, in agreement with an increase in methylation at two positions, 28S-Cm2861 and 28S-Gm2876, the upregulated expression of the corresponding guide SNORD50A was observed. In SH-SY5Y FUS KO proliferating and differentiated cells, hypermethylation of 18S-Um354 and 18S-Cm1272 was paralleled by elevated levels of corresponding guide snoRNAs, SNORD90 and SNORD66. Furthermore, an increase of modification at 18S-Gm436 was accompanied by a higher abundance of SNORD100 in differentiated SH-SY5Y FUS KO cells compared to WT cells. In SH-SY5Y cells expressing FUS R495X mutant, an increase in the methylation status of 18S-Cm1440 was paralleled by a higher level of the guide SNORD125, while hypomethylation of 28S-Am1323 was paralleled by the downregulation of SNORD126. Furthermore, changes in the methylation pattern of 28S-Cm1881, 28S-Am2401, 28S-Gm2876, 28S-Am3867, and 28S-Cm4456 were accompanied by consistent changes in the expression levels of corresponding snoRNAs: SNORD48, SNORD68, SNORD50A/SNORD50B, SNORD92, SNORD49A/SNORD49B in SH-SY5Y FUS KO differentiated cells, as evidenced by small RNA-seq data (Table 1, Fig. 3A, C). These results suggest that several changes in the methylation status of rRNAs observed in FUS KO and R495X cells can result directly from differential expression of particular snoRNAs affected by FUS (Fig. 4).

**Fig. 3.**
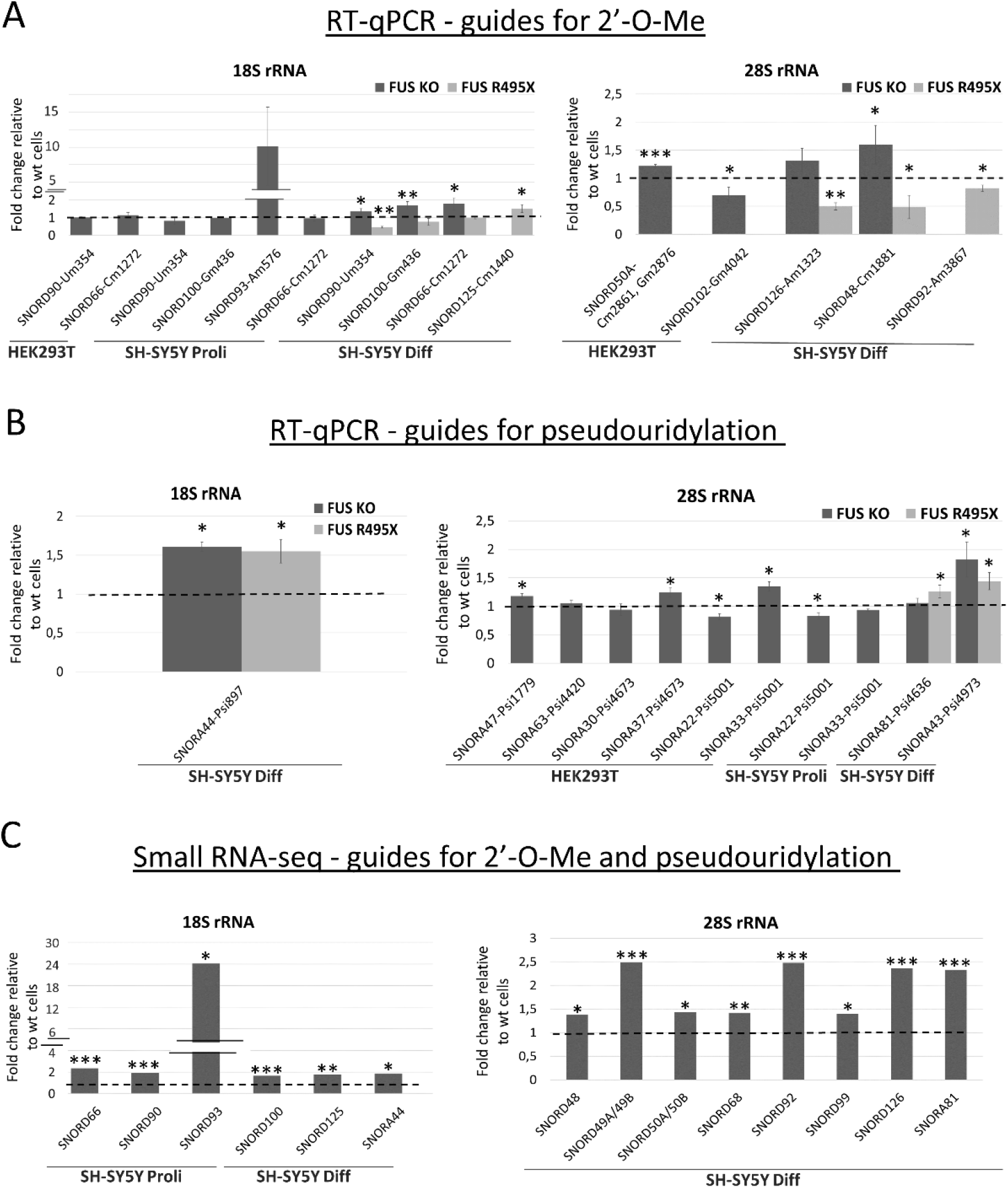
RT-qPCR analysis of relative expression levels of selected C/D box snoRNAs that guide methylation of residues in 18S rRNA and 28S rRNA (A); selected box H/ACA snoRNAs that guide pseudouridylation of residues in 18S rRNA and 28S rRNA (B). HEK293T FUS KO cells, proliferating (Proli), differentiated (Diff) SH-SY5Y FUS KO cells, and FUS R495X cells were compared to WT cells, respectively. Small RNAseq results of relative expression levels of selected differentially expressed snoRNAs that guide methylation and pseudouridylation of residues in 18S rRNA and 28S rRNA (C) in SH-SY5Y proliferating and differentiated cells. Error bars represent the SD of three biological replicates (N=3). P-values were calculated using Student’s t-test, and the statistical significance is defined as follows: *P ≤ 0.05; **P ≤ 0.01; ***P ≤ 0.001.

**Fig. 4.**
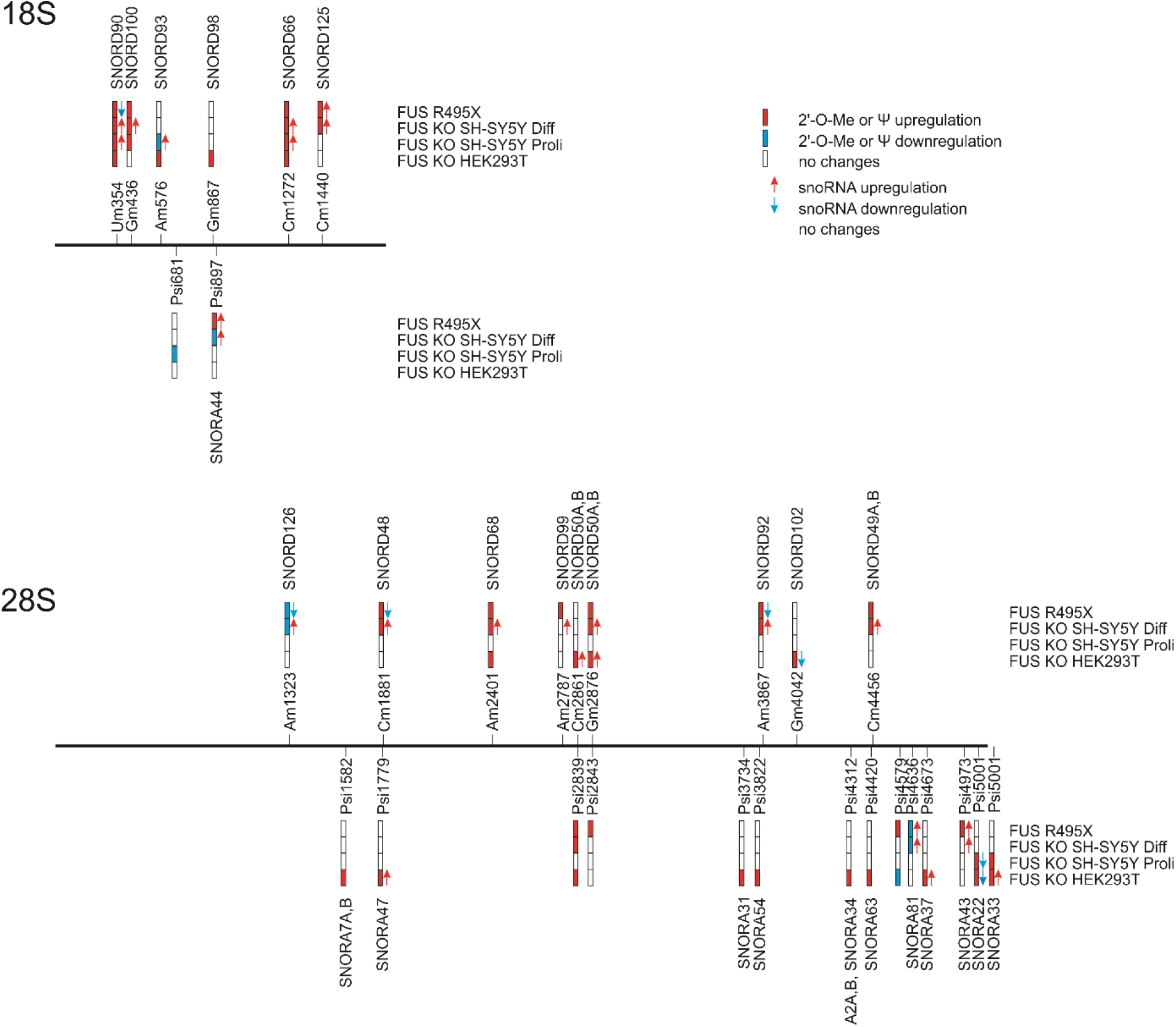
Scheme representing positions in 18S and 28S rRNAs with sites of methylation and pseudouridylation differentially modified in FUS KO and FUS R495X cells in comparison to wild-type cells. Red, blue and white bars indicate upregulation, downregulation or no changes, respectively. snoRNAs responsible for modifications are located above corresponding sites, red and blue arrows indicate snoRNA upregulation or downregulation, respectively, confirmed by small RNA-seq or RT-qPCR.

Moreover, to identify new 2’-O-Me sites in the rRNA that can be dependent on FUS, we used snoSCAN (40) tool with the snoRNA sequence fasta file from snoDB (41) and 18S and 28S rRNAs as target sequence. Interestingly, we were able to identify some putative 2’-O-Me sites in 18S rRNA (Cm951, Gm995, Cm1000, Am1034, Gm1639) and one site in 28S rRNA (Cm2075), all with low methylation scores and differential methylation patterns in FUS KO and FUS R495X compared to WT cells (Table 2, Supplementary Table S3). Furthermore, examining snoRNAs abundance using RT-qPCR and small RNA-seq data showed some correlations between methylation status and the expression of their corresponding snoRNAs in cells with FUS disturbances compared to WT cells (Table 2, Supplementary Fig. S4). The lack of methylation at 18S-Cm1000 in SH-SY5Y FUS KO proliferating cells was paralleled by the decreased expression level of SNORD44, as evidenced by RT-qPCR analysis. In the same cell line, 28S-Cm2075 was unmethylated in FUS KO cells, and this was accompanied by a lower level of SNORD114-14, as shown by small RNA-seq data. In SH-SY5Y differentiated cells, FUS depletion resulted in a 15% reduction in methylation score at 28S-Cm2075 and a consistent decrease in SNORD114-14 compared to WT cells, as evidenced by small RNA-seq data. Similarly, this position is hypomethylated in SH-SY5Y FUS R495X cells, followed by downregulation of SNORD114-14. Interestingly, SNORD114 family is considered an orphan snoRNA family, with no target determined. Another putative position, 18S-Cm951, was hypomethylated in SH-SY5Y FUS KO and FUS R495X differentiated cells, however, the predicted guide SNORD114-3 was downregulated only in FUS R495X cells. In SH-SY5Y FUS KO differentiated cells, a higher methylation score at 18S-Gm995 was paralleled by elevated expression of SNORD53, according to small RNA-seq data (Table 2, Supplementary Fig. S4). However, to confirm these putative positions further studies have to be performed.

**Table 2.** Putative 2’-O-methylation positions in rRNAs as identified by Snoscan (v.1.0).

| rRNA Subunit | Position and residue type | HEK293T WT | HEK293T FUS KO | C/D box snoRNA involved in 2'-O-methylation |
| --- | --- | --- | --- | --- |
|  |  | Av.MethScore | Av.MethScore |  |
| 18S | Cm951 | 0.34 | 0.01 ↓ | SNORD114-3 |
| 18S | Cm1000 | 0.29 | 0.14 ↓ | SNORD44 |
| 18S | Gm1639 | 0.62 | 0.49 ↓ | SNORD81 |
| 28S | Cm2075 | 0.21 | 0.04 ↓ | SNORD114-14 |

| rRNA Subunit | Position and residue type | SH-SY5Y WT Proli | SH-SY5Y FUS KO Proli | C/D box snoRNA involved in 2'-O-methylation |
| --- | --- | --- | --- | --- |
|  |  | Av.MethScore | Av.MethScore |  |
| 18S | Gm995 | 0.13 | 0.24 ↑ | SNORD53 |
| 18S | Cm1000 | 0.10 | 0.000 ↓ | SNORD44 |
| 18S | Am1034 | 0.12 | 0.30 ↑ | SNORD59A |
| 18S | Gm1639 | 0.56 | 0.42 ↓ | SNORD81 |
| 28S | Cm2075 | 0.20 | 0.000 ↓ | SNORD114-14 |

| rRNA Subunit | Position and residue type | SH-SY5Y WT Diff | SH-SY5Y FUS KO Diff | SH-SY5Y FUS R495X Diff | C/D box snoRNA involved in 2'-O-methylation |
| --- | --- | --- | --- | --- | --- |
|  |  | Av.MethScore | Av.MethScore | Av.MethScore |  |
| 18S | Cm951 | 0.36 | 0.13 ↓ | 0.08 ↓ | SNORD114-3 |
| 18S | Gm995 | 0.17 | 0.27 ↑ | 0.25 ↑ | SNORD53 |
| 18S | Gm1639 | 0.70 | 0.31 ↓ | 0.33 ↓ | SNORD81 |
| 28S | Cm2075 | 0.16 | 0.01 ↓ | 0.000 ↓ | SNORD114-14 |

The MethScores represent the average of three biological replicates (N=3) and describe a fraction of molecules methylated at a given position. Arrows depict the direction of modification changes in FUS KO and FUS R495X cells relative WT cells.

### FUS depletion and FUS R495X mutation affect pseudouridylation levels at specific positions in 18S and 28S rRNAs

To map and quantify the pseudouridine content in FUS KO and FUS R495X cells, we applied the HydraPsiSeq approach (30). This allowed us to profile the pseudouridylation level of annotated 44 sites in 18S rRNA, 61 sites in 28S rRNA, and two sites in 5.8S rRNA (Supplementary Table S4). Like the 2’-O-Me signatures, most pseudouridine sites were resistant to FUS depletion or mutation and exhibited near-saturated modification levels (Supplementary Table S4, Supplementary Fig. S3B). Interestingly, most of the significant variations in pseudouridylation between FUS KO or FUS R495X and WT cells (PsiScore difference > 0.05) refer to 28S rRNA, and similarly to methylation status, in most cases increased pseudouridine level in FUS knockout and mutant cells was observed (Table 1, Fig. 2B, Fig. 4). For example, the pseudouridylation status at site 28S-Psi5001 was significantly higher in HEK293T FUS KO and neuroblastoma FUS KO proliferating cells, while pseudouridylation status at site 28S-Psi2839 was increased in HEK293T FUS KO and neuroblastoma FUS KO and FUS R495X differentiated cells. In turn, pseudouridylation at position 28S-Psi4579 was lower in HEK293T FUS KO cells but higher in FUS R495X cells, compared to WT, with no changes in neuroblastoma FUS KO cells. This suggests that FUS can act differently depending on the cell line. Interestingly, position 18S-Psi897 was differentially modified in neuroblastoma differentiated cells, depending on whether FUS was depleted or only mutated (Table 1, Fig. 2B, Fig. 4).

In the next step, using previously described small RNA-seq data and RT-qPCR approach, we checked whether changes in modification level at specific positions might be accompanied by differential expression of corresponding guide snoRNAs in both FUS KO and FUS R495X mutant compared to WT cells (Supplementary Table 1, Fig. 3B, C, Fig. 4). In SH-SY5Y FUS R495X differentiated cells, increased pseudouridylation at 18S-Psi897 was accompanied by increased expression of the corresponding SNORA44. In HEK293T FUS KO cells, increased pseudouridylation at sites 28S-Psi1779, 28S-Psi4673, and 28S-Psi5001 was paralleled by elevated levels of the corresponding snoRNAs: SNORA47, SNORA37, and SNORA33, respectively. Interestingly, modifications at 28S-Psi4673 and 28S-Psi5001 are guided by two snoRNAs, but only one of them responds to FUS depletion (Table 1A, C, Fig. 3B, C, Fig. 4). Like in the 2’-O-methylation profile, in these particular examples we can assume that changes in the pseudouridylation status of rRNAs could be a direct consequence of the differential expression of specific snoRNAs affected by FUS. However, in other cases, the mechanisms which regulate post-transcriptional modifications of ribosomes are still elusive.

Next, we decided to elucidate the putative mechanism by which FUS can affect snoRNA levels and further modulate rRNA modifications as a secondary effect. As snoRNA genes are mostly localized within introns of other genes, so-called host genes, first we tested the expression of snoRNA host genes in cells with FUS depletion. As shown in Supplementary Fig. S5A, in HEK293T FUS KO cells the levels of selected host genes were generally upregulated, which only partially correspond to the elevated level of their hosted snoRNAs in this cell line. Further, by RNA immunoprecipitation (RIP) experiment we revealed that FUS can directly bind to both snoRNAs (Supplementary Fig. S5B, C) and selected snoRNA host gene transcripts (Supplementary Fig. S5D), however in this assay we couldn’t distinguish whether FUS can bind to mature snoRNA separately. Moreover, analysis of fibrillarin, NOP56, and dyskerin, which form snoRNP complexes, did not show spectacular changes in their expression in HEK293T and neuroblastoma cell lines with FUS knockout or mutation. Fibrillarin level slightly increased in SH-SY5Y cells, at both mRNA and protein level, expression of NOP56 protein was minimally induced in SH-SY5Y differentiated cells, whereas the expression of dyskerin was not altered in any cells with FUS changes (Supplementary Fig. S6).

### Mapping of FUS-dependent 2’-O-Me and pseudouridylation sites on the SSU and LSU rRNA subunits and 80S ribosome

It has been recently established that sites with dynamic 2’-O-Me or pseudouridylation scores can be responsible for the generation of ribosome heterogeneity (22,30,42–44). Even a single conserved 2’-O-Me site, for example, 18S-Cm174, can influence the translation of a specific set of mRNAs, highlighting the importance of individual sites of post-transcriptional modifications (25). We used PDB file 4UG0 in Pymol 2.0 to get LSU (28S, 5.8S, 5S rRNAs) and SSU (18S rRNA) structures (45) and map 2’-O-Me and pseudouridylation sites that were significantly changed in rRNA subunits in FUS KO and FUS R495X cells, followed by combining these subunits to reveal whether any of these sites are present within functionally conserved domains like E-site, DCS (decoding site), and PTC (peptidyl transferase centre) (Fig. 5). We identified that Um354 and Cm1440 sites within the 18S rRNA map to the outer periphery. These sites may be accessible for molecular interactions with ribosomal proteins, as opposed to the sites within PTC or DCS that are highly inaccessible (24). Remarkably, 18S-Cm1272 site which exhibited higher modification level in all FUS KO as well as in FUS R495X mutant cells, is located close to DCS. Usually, posttranscriptional modifications (PTMs), like 2’-O-Me and pseudouridylation, within the functionally important regions are very stable, however increase in 18S-Cm1272 2’-O-Me could indicate that this 2’-O-Me site within the DCS is sensitive to FUS. Another 2’-O-Me site, 18S-Cm1440, is present in the helix39-helix39ES (Expansion Segment), which is reported to be highly irregular and solvent accessible (Fig. 5) (24).

**Fig. 5.**
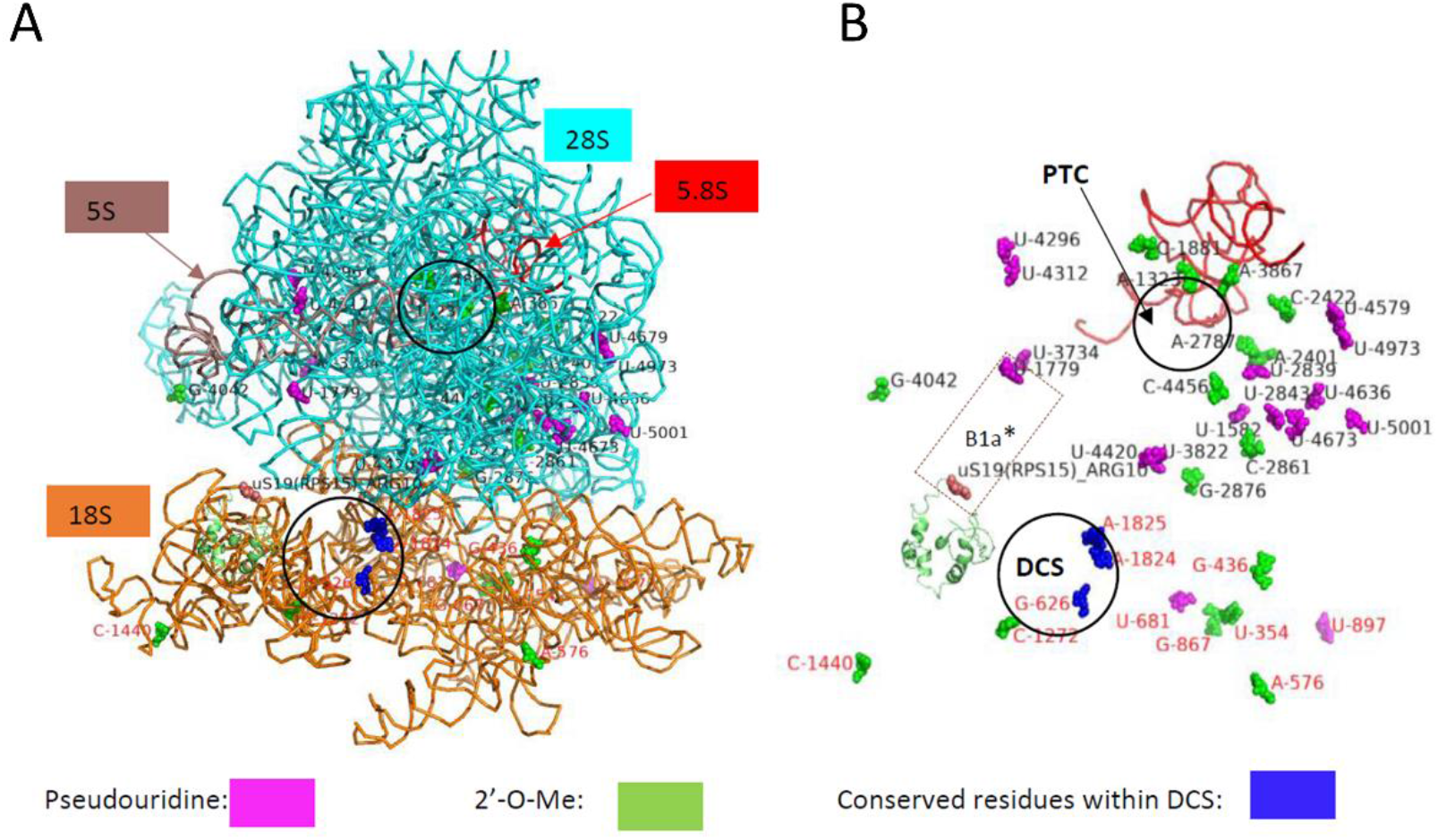
Visualization of the 2’-O-Me and pseudouridylation sites with the significant differences in FUS KO and FUS R495X cells on the LSU (28S, 5.8S, and 5S rRNA) and SSU (18S rRNA). Ribbon representation of PDB:4UG0 file. All the rRNAs are denoted by a unique colored ribbon and labelled in the same color (A). Representation of only modified residues, green for 2’-O-Me and magenta for pseudouridylation with only 5.8S rRNA in a red ribbon form to show structure orientation. uS19 is represented as a green color cartoon, with ARG10 denoted as brown-colored spheres. *B1a is an inter-subunit bridge formed by the interaction between ARG10 of uS19 and Psi-1779. DCS, decoding center (SSU), PTC, peptidyl transferase center (LSU) (B).

In the case of pseudouridylation, all but two significantly changed sites are within 28S rRNA (Table 1, Fig. 4). 28S-Psi1779 is of particular interest as the interaction between 28S-Psi1779 and uS19 (ribosomal protein S15-RPS15) Arginine 10 forms B1a inter-subunit bridge (45) (Fig. 5B). This site is hyperpseudouridylated in HEK293T FUS KO cells, and the corresponding guide RNA, SNORA47, is also upregulated in these cells (Fig. 4).

To estimate the effect of alterations in rRNA modifications on global translation efficiency we performed surface sensing of translation (SUnSET) assay. As shown in Supplementary Fig. S7, no changes were observed in HEK293T FUS KO cells, proliferating and differentiated SH-SY5Y FUS KO cells as well as in FUS R495X mutant cells, in comparison to wild-type cells. As observed in the latest studies using CRISPR-Cas9 method to knockout specific snoRNA which resulted in absence of corresponding modification on rRNA, we speculate that instead of global changes, translation of a subset of mRNAs can be specifically hampered or even translational fidelity is compromised (25,32).

### 2’-O-methylation and pseudouridylation levels of U snRNAs vary within cell lines and cell state

Small Cajal body-specific RNAs are responsible for site-specific 2’-O-methylation and pseudouridylation of snRNAs. Recent studies suggests the role of another RNA binding protein involved in ALS, TDP-43 (15,46), in regulating site-specific 2’-O-methylation of U1 and U2 snRNAs, by controlling the localization of a subset of C/D box scaRNAs (47). FUS is involved in major and minor intron splicing by its direct association with U1, U2, U11, and U12 snRNAs as well as in replication-dependent histone gene expression by interaction with U7 snRNA (9,16,36,48). These interactions are disturbed by ALS-associated mutations in FUS (16,37,49). Our small RNA-seq results confirmed that some C/D box and H/ACA box scaRNAs are differentially expressed in FUS KO cells (Supplementary Table S1). Next, to identify the effect of the altered scaRNAs expression on snRNAs, we checked the MethScores and PsiScores for known positions on snRNAs. Our results from RiboMeth-seq indicate that most of the snRNA positions are fully methylated in both proliferating and differentiated SH-SY5Y cells and we couldn’t observe any significant changes in 2’-O-Me levels in snRNAs in FUS R495X cells as well (Fig. 6A, Supplementary Table S2). However, we noted that cytosine at position Cm8 on U4 snRNA (RNU4-Cm8) is differentially methylated in HEK293T cells compared to neuroblastoma cells. Unfortunately, we could not detect known 2’-O-Me sites at the 5’ end of U1, U2, U4, and U5 snRNAs; RiboMethSeq is not appropriate to measure these terminal methylations.

**Fig. 6.**
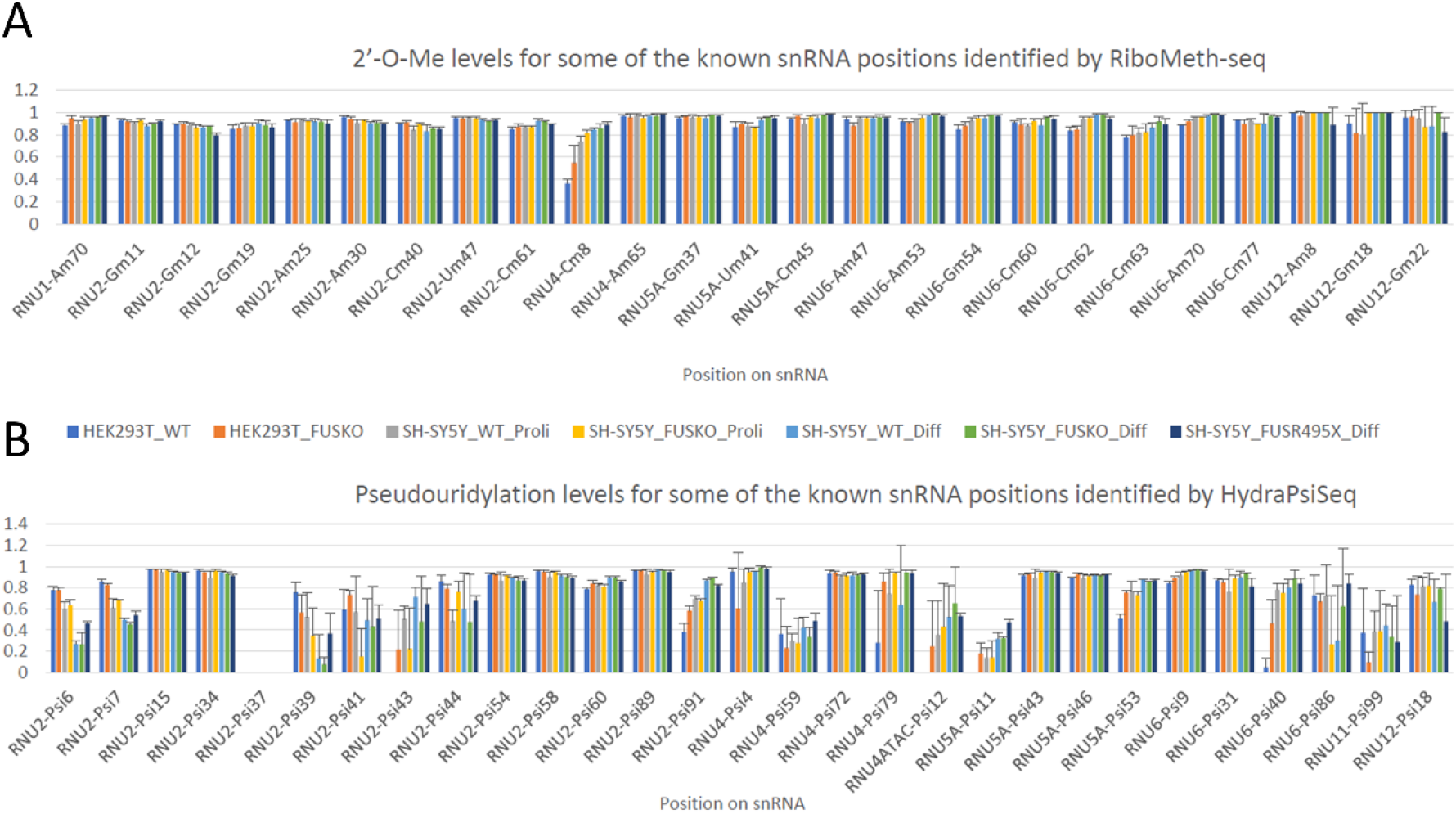
Distribution and extent of modifications of 2’-O-Me (A) and pseudouridylation (B) of residues in snRNAs. Bars represent the average of MethScore and PsiScore of three biological replicates (N=3) from each cell line. Error bars represent the SD (standard deviation) between three biological replicates.

By analyzing pseudouridylation levels of snRNAs, we found an increase of this modification, for example, at position Psi6 of U2 snRNA (RNU2-Psi6) in SH-SY5Y R495X cells (Fig. 6B, Supplementary Table S4). As reported by Yu et al., PTMs of the 5’ end region of U2 snRNA are necessary to form functional U2 snRNP and pre-mRNA splicing (27). Moreover, we observed increased pseudouridylation at position Psi91 (RNU2-Psi91) in HEK293T FUS KO cells. Noticeable differences were observed within SH-SY5Y WT cells in proliferating and differentiated conditions, specifically for sites RNU2-Psi6, RNU2-Psi7, and RNU2-Psi91 (Fig. 6B). It is essential to mention that we observed variability in PsiScore within replicates, resulting from lower sequencing depth; because of that we also could not precisely measure all the known pseudouridylation positions in snRNAs.

## DISCUSSION

### 2’-O-Me and pseudouridylation changes in FUS KO and FUS R495X cells and their implications in ALS

Our study highlights that depletion and mutation of FUS, an RNA binding protein, can result in differential post-transcriptional modifications of rRNAs. Further, the changes observed in 2’-O-Me and pseudouridylation levels in rRNAs could indicate changes in ribosome fidelity although we did not observe significant changes in global translation efficiency in FUS KO and FUS R495X mutant cells (Supplementary Fig. S7). Moreover, we also report that ALS-associated FUS mutations can result in changes at the level of post-transcriptional modifications of ribosomal RNAs.

Few previous studies reported that snoRNAs are enriched in sporadic ALS patient samples compared to the healthy controls (39,50). We observed that snoRNAs are mostly upregulated in SH-SY5Y FUS KO cells differentiated with retinoic acid (Fig. 1B, Fig. 4, Supplementary Table S1). Even though these studies do not specifically associate FUS with snoRNA levels, there is an indication that some snoRNAs, such as SNORD48, SNORD90 and SNORD100, which are upregulated in SH-SY5Y FUS KO cells, are also enriched in sporadic ALS patients (39,50). However, how FUS controls the expression of snoRNAs is still not well understood. It is known from previous work that wild-type FUS explicitly binds introns within pre-mRNAs (46); interestingly, most of the snoRNAs are located in the introns of protein-coding or non-coding RNA genes. As we reported here, the changes in the level of host gene transcripts and their encoding snoRNAs do not always correspond to each other (Supplementary Fig. S5A), therefore in these cases we may exclude that FUS regulates snoRNA level indirectly by the regulation of their host gene transcription or splicing. Nevertheless, in cases where similar changes of both host genes transcripts and snoRNAs were observed in FUS KO cells, FUS might affect snoRNA maturation instead of acting on mature snoRNAs. One should also keep in mind that many host genes encode more than one snoRNA genes (such as RPS12, CCNB1IP1, SNHG32, SNHG12, WDR43, EIF4A2) and thus host gene transcript levels do not always mirror the level of expression of all their snoRNAs. Unfortunately, our RIP approach prevented distinguishing mature snoRNAs from their host gene transcripts in the FUS-immunoprecipitated fractions, leaving this question still open (Supplementary Fig. S5B-D). Additionally, we analyzed publicly available CLIPseq datasets hosted on POSTAR3 database (51). Interestingly, we observed that FUS interacts with some snoRNAs, however, in many cases the binding positions overlap with positions identified for their host gene transcripts, precluding interpretation of which mechanisms, host gene transcript level or mature snoRNA level, is directly affected by FUS. Moreover, our analysis showed no or little changes in the expression of fibrillarin, dyskerin and NOP56 in cells with FUS alterations, excluding the hypothesis that changes in posttranscriptional modifications of rRNAs and snRNAs results from disrupted assembly of snoRNP complexes. Summarizing, the mechanism by which FUS can affect snoRNA levels and therefore indirectly modify rRNA modifications, is still not elucidated.

As reported by multiple previous studies, fractionally modified 2’-O-Me and pseudouridylation sites are responsive to changes, such as stress, differentiation, and knockdown or mutation of vital snoRNP factors, including fibrillarin, dyskerin, and NOP10 (23,42,44,52,53). H/ACA box snoRNA levels are regulated during stem cell differentiation (52), which could also indicate a downstream effect on the level of pseudouridylation within rRNA and snRNAs guided by this class of snoRNAs. By RiboMeth-seq we were able to identify all known 2’-O-Me sites within ribosomal RNAs. Even though most of these sites were almost fully methylated, we observed a general increase in 2’-O-Me levels in FUS KO cells (Supplementary Table S2) which coincides with our observation that snoRNAs are enriched upon FUS depletion (Supplementary Table S1). It is important to mention that we observed only a small proportion of modified sites that actually correlate with the expression of corresponding guide snoRNAs (Table 1, Fig. 3, Fig. 4). Our observation is consistent with previous reports suggesting that apart from the snoRNA levels, possibly there are additional factors involved in the regulation of post-transcriptional modifications of ribosomal RNAs (23,24,42). Interestingly, Um354 and Gm436 sites within 18S rRNA showed increased 2’-O-Me; these sites are located in the 5’ domain of helix7 and helix13. An earlier study by Krogh et al. suggests that an increase in 2’-O-Me level at these positions may reduce cell growth due to decreased ribosomal efficiency (24); likewise, we observed almost saturated level of modification at these sites in FUS R495X cells. Although in our analysis we did not observe changes in global translation efficiency in cells with FUS R495X mutation (Supplementary Fig. S7), the hyper-modification at these sites can contribute to the global suppression of protein synthesis observed in mutant FUS-associated ALS patients, which so far was linked to increased binding of ALS-FUS mutant proteins to the polyribosome fraction as well as sequestration of proteins involved in translation into cytoplasmic mutant FUS inclusions (3,54). Moreover, we do not rule out the possibility that the modification changes observed in cells with FUS disturbances can affect translation of only specific pool of mRNA molecules and also translational fidelity, which is impossible to detect by measuring global translation.

Furthermore, we identified scaRNAs that are differentially expressed in FUS KO cells and some pseudouridylation sites in snRNAs that were altered in these cells and in FUS R495X mutant (Supplementary Table S1, Fig. 6B) which could partially explain RNA splicing defects reported in FUS depleted cells and caused by ALS-associated FUS mutations (33–35).

Interestingly, apart from the snoRNAs that act as guide RNAs for ribosomal and snRNA modifications, we also identified some snoRNAs from genetically-imprinted, orphan, C/D box snoRNA families, like SNORD113, SNORD114, and SNORD116, to be differentially expressed in SH-SY5Y FUS KO cells with the SNORD114 family member potentially targeting 28S-Cm2075 (Table 2, Supplementary Fig. S4). Several patients with Prader-Willi Syndrome (PWS) phenotype have microdeletions within SNORD115 and SNORD116 clusters (55,56). SNORD113 and SNORD114 families are encoded within maternally-imprinted host gene MEG8 and are enriched in acute leukemia, suggesting a possible role in oncogenesis (57).

### Differences in 2’-O-Me and pseudouridylation levels within cell lines and cell state

As previously shown, the methylation level for stably modified residues within rRNAs is relatively high, with MethScore > 0.9 (44). Similarly, in our analysis, 72/109 (66%) positions are highly modified with MethScore > 0.9. For some positions we observed a significant difference in methylation score between WT cell lines that differed by the threshold of 10% (Fig. 2A, Supplementary Fig. S3A, Supplementary Table S2). Interestingly, methylation pattern at three positions, 18S-Am576, 5.8S-Um14, and 28S-Gm2876, vary significantly between HEK293T and neuroblastoma proliferating cells. In addition, 28S-Gm2876 methylation status was consistently influenced by FUS disturbances, as evidenced by the increased 2’-O-Me level in HEK293T FUS KO cells and differentiated SH-SY5Y FUS KO and FUS R495X cells. 18S-Gm436 differs only between HEK293T and SH-SY5Y in proliferating conditions, and its methylation level was also sensitive to FUS depletion. Moreover, we identified a few sites that vary between SH-SY5Y cells in proliferating and retinoic acid-mediated differentiated states, specifically 18S-Um354, 18S-Gm436, 18S-Cm1272, 18S-Cm1440, 28S-Am1323, and 5.8S-Um14. Interestingly, all these sites but 5.8S-Um14 appeared to be sensitive to FUS depletion in at least one of the conditions (Fig. 2A, Supplementary Fig. S3A, Supplementary Table S2).

Comparison of HydraPsiSeq profiles obtained for HEK293T WT cells and SH-SY5Y WT proliferating and differentiated cells showed that a subset of detected pseudouridylation sites exhibited considerable variations in PsiScore levels (cut off 10%). Several positions displayed distinctly different PsiScore levels between HEK293T and neuroblastoma cells, such as 28S-Psi4579, 18S-Psi681, 18S-Psi1046, 28S-Psi1536, 18S-Psi296, 28S-Psi3884. Surprisingly, we identified only one site, 28S-Psi1536, with significant hypermodification in SH-SY5Y differentiated cells compared to those in proliferating state (Fig. 2B, Supplementary Fig. S3B, Supplementary Table S4).

In summary, our analysis shows that methylation and pseudouridylation profiles of human rRNAs may vary between different cell lines and due to the differentiation process. Furthermore, we hypothesize that a subset of fractionally modified 2’-O-Me and pseudouridylation sites on rRNAs might decide about the change from proliferating to differentiated cell state. These results are consistent with previously published data and support the notion that different cell types can produce heterogeneous, differentially modified populations of ribosomes with specialized translation properties (24,25,44). To conclude, our study reports increased expression of a subset of snoRNAs in FUS depleted and ALS-associated FUS mutant cells. The increase in snoRNA expression correlates with higher modification at the corresponding positions on rRNAs. We thereby report a novel mechanism which might be disturbed in ALS and contribute to disease pathology.

## MATERIAL AND METHODS

### Cell culture and differentiation

SH-SY5Y and HEK293T cells were grown in Dulbecco’s modified Eagle’s medium containing L-glutamine and 4.5 g*/*L glucose (DMEM; Lonza) and supplemented with 10% fetal calf serum (Gibco) and antibiotics (100 U*/*ml penicillin, 100 µg*/*ml streptomycin (Sigma-Aldrich)) at 37°C in a humidified atmosphere containing 5% CO_2_. SH-SY5Y cells with FUS KO were prepared as described previously (34,58). HEK293T cells with FUS KO were generated according to (59), using two SpCas9-2A-Puro (RX459) plasmids coding for sgRNAs: 5’-GCGTCGGTACTCAGCGGTGT-3’, 5’-GCCCGGGCAGGGCTATTCCC-3’ targeting exon 1 and exon 3 of the *FUS* gene, respectively. This strategy resulted in the deletion of the genomic fragment of 2360 bp. Knockout efficiency was checked by DNA sequencing and Western blot followed by immunodetection with anti-FUS antibody (Supplementary Fig. S8A). Exon 14 of the *FUS* gene was targeted to introduce the R495X mutation in SH-SY5Y cells using the pCRISPR-EF1a-SpCas9-R495X plasmid coding for the sgRNA targeting the sequence: 5′-GGGACCGTGGAGGCTTCCGA-3′, following the strategy described in (49). The plasmids used to generate SH-SY5Y FUS R495X cells were obtained from the Marc-David Ruepp group and used in two transfection approaches. In the first approach, 8×10^5^ cells were seeded in two wells of a six-well plate one day before transfection and transfected using VIROMER® RED (Lipocalyx) according to the manufacturer’s instructions. Each well was transfected with 3 µg of plasmid DNA, including 120 ng of pRR-Puro-R495X, 1950 ng of pCRISPR-EF1a-SpCas9-R495X, and 920 ng of R495X donor plasmid for HDR. In the second approach, SH-SY5Y cells seeded in two wells of a 24-well plate were transfected with Lipofectamine 2000 according to the manufacturer’s instructions with a total amount of 500 ng of plasmid DNA, including 20 ng of pRR-Puro-R495X, 240 ng of pCRISPR-EF1a-SpCas9-R495X and 240 ng of R495X donor plasmid for HDR. 24 hours after transfection puromycin selection was applied to each well of two different transfection strategies at a concentration of 2 µg/ml (54 h) or 1 µg/ml (5 days), respectively. After selection, the cells from both transfection approaches were grown in culture medium without antibiotics for 7 more days and then pooled; clonal cell lines were isolated by serial dilution. Subsequently, colonies grown from single cells were picked, and gDNA was isolated for clone screening using the ExtractMe Genomic DNA Kit (Blirt). The R495X genomic locus was amplified from the genomic DNA using DreamTaq DNA polymerase (Thermo Scientific) (Supplementary Table S5). PCR products were purified using the GeneJET Gel Extraction and DNA Cleanup Micro Kit (Thermo Scientific) and sequenced (Supplementary Fig. S8B). The expression and partial mislocalization of mutated FUS in cytoplasmic aggregates in SH-SY5Y differentiated cells was tested by Western blot and immunodetection and by immunofluorescence with anti-FUS antibody, respectively (Supplementary Fig. S8C). In all cell lines with FUS knockout and mutations, the level of EWSR1 and TAF15 was also tested showing either no changes or gentle upregulation of both proteins (Supplementary Fig. S8D).

For cellular differentiation, SH-SY5Y wild-type, FUS KO, and FUS R495X cells were subjected to retinoic acid (RA) treatment for 10 days to transform them into neuron-like cells (16,60). All-trans RA (USP, Tretinoin, 1674004) at a final concentration of 75 µM was added to DMEM with 10% FBS. The media, along with the RA, were replaced every 3 days. The differentiation efficiency was analyzed as described in (16).

### RNA isolation, cDNA preparation, and qPCR

Total RNA was isolated using the Quick-RNA MiniPrep Kit (Zymo Research, R1055) according to the manufacturer’s protocol. First-strand cDNAs were synthesized in 25 μl reactions with 1 μg of RNA using 200 ng of random hexamer primers and 200 U Superscript III reverse transcriptase (Thermo Scientific), according to the manufacturer’s protocol. For qPCR, 1 µl of 5× diluted cDNA template, 0.2 µM primer mix (forward + reverse), and 5 µl of SYBR Green PCR master mix (Applied Biosystems) were added in a 10 µl reaction with the following conditions: denaturation for 10 min at 95°C, followed by 40 cycles of 95°C for 15 s and 60°C for 1 min (Applied Biosystems QuantStudio 7 Flex). Primers used for qPCR reactions are listed in Supplementary Table S5. The statistical significance of the qPCR results was determined by Student’s t-test.

### Western blot analysis and immunofluorescence

Protein extraction was performed using gentle hypotonic lysis buffer (10 mM Tris pH 7.5, 10 mM NaCl, 2 mM EDTA, 0.5% Triton-X-100) supplemented with 1x cOmplete protease inhibitor (Roche). After 30 min incubation on ice, the NaCl concentration was adjusted to 150 mM, followed by another 5 min incubation on ice. The lysate was cleared by centrifugation at 16 000xg for 15 min at 4 °C. Western blot analyses were performed according to the already published protocol from our laboratory (16). Briefly, 30 μg of proteins were separated by SDS–polyacrylamide gel electrophoresis, transferred to polyvinylidene difluoride (PVDF) membranes (Millipore), blocking was done with 5% skim milk diluted in PBS or TBS with 0.1 % Tween-20 (PBS-T or TBS-T), and then incubated for 1.5-2h h at room temperature (RT) with anti-FUS (Santa Cruz, 4H11, 1:3000 dilution in PBS-T), home-made polyclonal rabbit anti-FUS antibody (9), anti-actin (MP Biomedicals, 691001, 1:25,000 dilution in PBS-T), anti-FBL (Santa Cruz, sc-25397, 1:500 dilution in TBS-T), anti-NOP56 (Santa Cruz, sc-23701, 1:500 dilution in TBS-T), anti-EWSR1 (Santa Cruz, sc-48404, 1:500 dilution in TBS-T) and anti-puromycin (Merck, MABE343, 1: 5000 dilution in TBS-T) primary antibodies. After washing the membrane with PBS-T for 3 times, it was incubated for 1 h at RT with species-specific horseradish peroxidase (HRP)-coupled secondary antibody (Santa Cruz, sc-2005, goat anti-mouse; Santa Cruz, sc-2357, mouse anti-rabbit; Santa Cruz, sc-2020, monkey anti-goat; 1:3000 dilution in PBS-T). 3 washes with PBS-T were performed as described earlier. The signal was detected using the enhanced chemiluminescence method (ECL, GE Healthcare). Immunofluorescence was performed according the protocol described in (16).

### RNA immunoprecipitation

For RNA immunoprecipitation, a total of 225 μg of protein extract isolated using the aforementioned hypotonic buffer (additionally supplemented with 100 U/ml RNasin Plus RNase Inhibitor (Promega) and 2 U/mL Turbo DNase (Ambion)) was incubated with Dynabeads Protein G conjugated with home-made polyclonal rabbit anti-FUS antibody for 2 h at 4°C. After three washes with PBS–T, the beads were resuspended in 0.5 ml TRIZOL and RNA was isolated according to the protocol described in (9). As input, 25 µg of protein extract was directly added to 0.5 ml of TRIZOL. cDNA was synthesized using random hexamer primers. As a negative control for RIP experiment, Dynabeads Protein G coupled to mouse IgG were used.

### SUnSET assay

For SUnSET assay, cells were treated with 1µM puromycin (Gibco) for 30 minutes according to the protocol described in (54).

### Small RNA sequencing library preparation and data analysis

500 ng of total RNA were used to prepare small RNA sequencing libraries using Small RNA-Seq Library Prep Kit (Lexogen, Vienna, Austria) following the manufacturer’s protocol. Library quality was analyzed with High Sensitivity DNA Kit on Agilent Bioanalyzer 2100, and quantification was done using Qubit dsDNA HS (High Sensitivity) Assay kit (Invitrogen™) with Qubit fluorometer. Quantified libraries were sent to the Lexogen sequencing facility (Vienna, Austria). Single-end read sequencing was carried out with 75 bp sequencing length on the Illumina NextSeq500 platform. All analyses were done in three biological replicates. The obtained FASTQ files were first subject to quality check with FastQC v0.11.4 (61). Then, fastx-toolkit/0.0.14 (62) was used for adapter removal and quality filtering of the FASTQ files. The adapter clipped reads were also aligned to ribosomal RNA sequences using Bowtie 2 (v2.2.3) (63), and the mapped reads were discarded. After that, mapping against the human genome (GRCh38 downloaded from Ensembl) was performed with Bowtie 2 (v2.2.3). The resulting SAM files were converted to BAM format using samtools (64) and were subsequently applied with an R package Rsubread featureCounts (65) to get raw read counts with the parameters *allowMultiOverlap=TRUE* and *countMultiMappingReads=TRUE*. A GTF file was created from Homo_sapiens.GRCh38.100.gtf (Ensembl) that consists of only snoRNAs, miRNAs, and snRNAs used in *featureCounts* as an annotation file. The obtained raw counts matrix was finally applied to identify differentially expressed genes (DEGs) using DESeq2 (66) with default settings and a threshold of 0.05 for adjusted P-value. Heatmaps were generated only with DESeq normalised snoRNA reads with gplots (67) from R. Volcano plots were generated using Enhanced Volcano R package (68).

### Library preparation and RiboMeth-seq (RMS)

The protocol was adapted from (29). In short, 5 μg of total RNA was used for each replicate, and alkaline degradation was performed at denaturing temperatures. 20-40 nt library fragments were gel-purified on a polyacrylamide-urea gel followed by ligation of 5’ and 3’ end linkers using tRNA ligase from *Arabidopsis thaliana*. 3’ adapter has a sequence barcode used as a primer site for sequencing. 2’ phosphates were removed in the subsequent purification step, and libraries were subjected to sequencing on the Ion Proton platform (Life Technologies). The sequencing depth for each biological replicate was between 80 and 100 million reads. Data analysis of RiboMeth-seq: The data analysis was performed as described previously (42). Briefly, a corrected human ribosomal DNA complete repeating unit was used (Genbank accession no. U13369) as a reference sequence (42). The libraries were generated from 20-40 nt fragments; hence, the ends of rRNA molecules were checked for ∼20 nt from one end. The MethScores mentioned in this study describe a fraction of molecules methylated at that position. The calculation is done by comparing number of 3’ and 5’ read ends at the inquired position to six flanking nucleotides on either sites (29). Heatmaps were generated with gplots (67) from R, representing the average of 3 biological replicates (N=3) from each cell line.

### RNA fragmentation and library preparation for HydraPsiSeq

Analysis of RNA pseudouridylation was performed according to previously published protocol (30). To summarize, 250 ng of RNA from each replicate was used for hydrazine treatment for 30-60 min on ice. To stop the reaction, ethanol precipitation with 0.3 M NaOAc, pH 5.2 and Glycoblue was performed, and incubated at −80°C for 30 min, followed by centrifugation. The collected pellet was washed with 80% ethanol and resuspended in 1 M aniline, pH 4.5, followed by incubation in the dark for 15 min at 60°C and then ethanol precipitation. Dephosphorylation of RNA at the 3’ end was done as described in (30). Library preparation was carried out using the NEBNext Small RNA Library kit (NEB, UK), following the manufacturer’s protocol. Library quality and quantity were assessed using a High Sensitivity DNA chip on an Agilent Bioanalyzer 2100 and a Qubit fluorometer (Invitrogen, USA). Multiplexed libraries were subjected to high-throughput sequencing using an Illumina NextSeq2000 instrument with single-read runs having 50 bp read length. HydraPsiSeq data analysis: Data analysis was performed as described in (30). In short, Trimmomatic v32.0 was used for adapter removal, and very short reads <8 nt were excluded. Bowtie 2 was used for the alignment of raw reads in end-to-end mode. In the next step, mapped and sorted .bam files were changed to bed format. Using a rolling window of 10 nucleotides, 5’ end counts were normalized to the local background. NormUcount was used as a protection Uscore value and transformed to U cleavage profiles by omitting values for other nucleotides. U profiles resulting from the previous step were used to calculate ScoreMEAN, A, B, and PsiScore as previously described (30). Three biological replicates were used for the analysis from each cell line, except for SH-SY5Y wild-type proliferating cells where only 2 replicates were considered for the final analysis.

### Statistical analysis

Paired Student’s t-test was used to calculate the statistical significance between samples from RT-qPCR, RiboMeth-seq and HydraPsiSeq. The statistical significance is represented as follows: *p<0.05; **p<0.01; ***p<0.001.

## Data availability

The small RNA-seq, RiboMeth-seq, and HydraPsiSeq data will be available at NCBI GEO. Supplementary Tables S1, S2 and S4 are available on request.

## ACKNOWLEDGEMENTS

We would like to thank Marc-David Ruepp’s group for the plasmids used for the generation of SH-SY5Y FUS R495X cell line and Anna Karlik for help with immunofluorescence experiment. This work was supported by the Polish National Science Centre [UMO-2018/30/E/NZ2/00295 to K.D.R.]; and the Initiative of Excellence-Research University project financed by the Polish Ministry of Science and Higher Education [ID-UB-003/13/UAM/006 to K.G. and ID-UB-038/04/NP/0008 to P.P.]. The data analysis was performed under the computational grant number 312 from the PCSS (Poznan Supercomputing and Networking Centre).

## REFERENCES

1. Tan, A.Y. and Manley, J.L. (2009) The TET family of proteins: functions and roles in disease. J. Mol. Cell. Biol., 1, 82–92.

2. Zinszner, H., Sok, J., Immanuel, D., Yin, Y. and Ron, D. (1997) TLS (FUS) binds RNA in vivo and engages in nucleo-cytoplasmic shuttling. J. Cell. Sci., 110 (Pt 15), 1741–1750.

3. Sévigny, M., Bourdeau Julien, I., Venkatasubramani, J.P., Hui, J.B., Dutchak, P.A. and Sephton, C.F. (2020) FUS contributes to mTOR-dependent inhibition of translation. J. Biol. Chem., 295, 18459–18473.

4. Perrotti, D., Bonatti, S., Trotta, R., Martinez, R., Skorski, T., Salomoni, P., Grassilli, E., Lozzo, R.V., Cooper, D.R. and Calabretta, B. (1998) TLS/FUS, a pro-oncogene involved in multiple chromosomal translocations, is a novel regulator of BCR/ABL-mediated leukemogenesis. EMBO J., 17, 4442–4455.

5. Baechtold, H., Kuroda, M., Sok, J., Ron, D., Lopez, B.S. and Akhmedov, A.T. (1999) Human 75-kDa DNA-pairing protein is identical to the pro-oncoprotein TLS/FUS and is able to promote D-loop formation. J. Biol. Chem., 274, 34337–34342.

6. Gardiner, M., Toth, R., Vandermoere, F., Morrice, N.A. and Rouse, J. (2008) Identification and characterization of FUS/TLS as a new target of ATM. Biochem. J., 415, 297–307.

7. Levone, B.R., Lenzken, S.C., Antonaci, M., Maiser, A., Rapp, A., Conte, F., Reber, S., Mechtersheimer, J., Ronchi, A.E., Mühlemann, O., et al. (2021) FUS-dependent liquid-liquid phase separation is important for DNA repair initiation. J. Cell. Biol., 220, e202008030.

8. Sama, R.R.K., Ward, C.L. and Bosco, D.A. (2014) Functions of FUS/TLS from DNA repair to stress response: implications for ALS. ASN Neuro, 6, 1759091414544472.

9. Raczynska, K.D., Ruepp, M.-D., Brzek, A., Reber, S., Romeo, V., Rindlisbacher, B., Heller, M., Szweykowska-Kulinska, Z., Jarmolowski, A. and Schümperli, D. (2015) FUS/TLS contributes to replication-dependent histone gene expression by interaction with U7 snRNPs and histone-specific transcription factors. Nucleic Acids Res., 43, 9711–9728.

10. Morlando, M., Dini Modigliani, S., Torrelli, G., Rosa, A., Di Carlo, V., Caffarelli, E. and Bozzoni, I. (2012) FUS stimulates microRNA biogenesis by facilitating co-transcriptional Drosha recruitment. EMBO J., 31, 4502–4510.

11. Zhang, T., Wu, Y.-C., Mullane, P., Ji, Y.J., Liu, H., He, L., Arora, A., Hwang, H.-Y., Alessi, A.F., Niaki, A.G., et al. (2018) FUS Regulates Activity of MicroRNA-Mediated Gene Silencing. Mol. Cell, 69, 787-801.e8.

12. Kwiatkowski, T.J., Bosco, D.A., Leclerc, A.L., Tamrazian, E., Vanderburg, C.R., Russ, C., Davis, A., Gilchrist, J., Kasarskis, E.J., Munsat, T., et al. (2009) Mutations in the FUS/TLS gene on chromosome 16 cause familial amyotrophic lateral sclerosis. Science, 323, 1205–1208.

13. Vance, C., Rogelj, B., Hortobágyi, T., De Vos, K.J., Nishimura, A.L., Sreedharan, J., Hu, X., Smith, B., Ruddy, D., Wright, P., et al. (2009) Mutations in FUS, an RNA processing protein, cause familial amyotrophic lateral sclerosis type 6. Science, 323, 1208–1211.

14. Dormann, D. and Haass, C. (2013) Fused in sarcoma (FUS): an oncogene goes awry in neurodegeneration. Mol. Cell. Neurosci., 56, 475–486.

15. Lagier-Tourenne, C., Polymenidou, M. and Cleveland, D.W. (2010) TDP-43 and FUS/TLS: emerging roles in RNA processing and neurodegeneration. Hum. Mol. Genet., 19, R46–64.

16. Gadgil, A., Walczak, A., Stępień, A., Mechtersheimer, J., Nishimura, A.L., Shaw, C.E., Ruepp, M.-D. and Raczyńska, K.D. (2021) ALS-linked FUS mutants affect the localization of U7 snRNP and replication-dependent histone gene expression in human cells. Sci. Rep., 11, 11868.

17. Ojha, S., Malla, S. and Lyons, S.M. (2020) snoRNPs: Functions in Ribosome Biogenesis. Biomolecules, 10, E783.

18. Bratkovič, T. and Rogelj, B. (2014) The many faces of small nucleolar RNAs. Biochim. Biophys. Acta, 1839, 438–443.

19. Filipowicz, W. and Pogacić, V. (2002) Biogenesis of small nucleolar ribonucleoproteins. Curr. Opin. Cell Biol., 14, 319–327.

20. Jaafar, M., Paraqindes, H., Gabut, M., Diaz, J.-J., Marcel, V. and Durand, S. (2021) 2’O-Ribose Methylation of Ribosomal RNAs: Natural Diversity in Living Organisms, Biological Processes, and Diseases. Cells, 10, 1948.

21. Penzo, M. and Montanaro, L. (2018) Turning Uridines around: Role of rRNA Pseudouridylation in Ribosome Biogenesis and Ribosomal Function. Biomolecules, 8, E38.

22. Jack, K., Bellodi, C., Landry, D.M., Niederer, R.O., Meskauskas, A., Musalgaonkar, S., Kopmar, N., Krasnykh, O., Dean, A.M., Thompson, S.R., et al. (2011) rRNA pseudouridylation defects affect ribosomal ligand binding and translational fidelity from yeast to human cells. Mol. Cell, 44, 660–666.

23. Sharma, S., Marchand, V., Motorin, Y. and Lafontaine, D.L.J. (2017) Identification of sites of 2’-O-methylation vulnerability in human ribosomal RNAs by systematic mapping. Sci. Rep., 7, 11490.

24. Krogh, N., Asmar, F., Côme, C., Munch-Petersen, H.F., Grønbæk, K. and Nielsen, H. (2020) Profiling of ribose methylations in ribosomal RNA from diffuse large B-cell lymphoma patients for evaluation of ribosomes as drug targets. NAR Cancer, 2, zcaa035.

25. Jansson, M.D., Häfner, S.J., Altinel, K., Tehler, D., Krogh, N., Jakobsen, E., Andersen, J.V., Andersen, K.L., Schoof, E.M., Ménard, P., et al. (2021) Regulation of translation by site-specific ribosomal RNA methylation. Nat. Struct. Mol. Biol., 28, 889–899.

26. Bohnsack, M.T. and Sloan, K.E. (2018) Modifications in small nuclear RNAs and their roles in spliceosome assembly and function. Biol. Chem., 399, 1265–1276.

27. Yu, Y.T., Shu, M.D. and Steitz, J.A. (1998) Modifications of U2 snRNA are required for snRNP assembly and pre-mRNA splicing. EMBO J., 17, 5783–5795.

28. Nagasawa, C.K., Kibiryeva, N., Marshall, J., O’Brien, J.E. and Bittel, D.C. (2020) scaRNA1 Levels Alter Pseudouridylation in Spliceosomal RNA U2 Affecting Alternative mRNA Splicing and Embryonic Development. Pediatr. Cardiol., 41, 341–349.

29. Birkedal, U., Christensen-Dalsgaard, M., Krogh, N., Sabarinathan, R., Gorodkin, J. and Nielsen, H. (2015) Profiling of ribose methylations in RNA by high-throughput sequencing. Angew. Chem. Int. Ed. Engl., 54, 451–455.

30. Marchand, V., Pichot, F., Neybecker, P., Ayadi, L., Bourguignon-Igel, V., Wacheul, L., Lafontaine, D.L.J., Pinzano, A., Helm, M. and Motorin, Y. (2020) HydraPsiSeq: a method for systematic and quantitative mapping of pseudouridines in RNA. Nucleic Acids Res., 48, e110.

31. Khoshnevis, S., Dreggors-Walker, R.E., Marchand, V., Motorin, Y. and Ghalei, H. (2022) Ribosomal RNA 2′-O-methylations regulate translation by impacting ribosome dynamics. Proc. Natl. Acad. Sci. U.S.A., 119, e2117334119.

32. Liang, J., Li, G., Liao, J., Huang, Z., Wen, J., Wang, Y., Chen, Z., Cai, G., Xu, W., Ding, Z., et al. (2022) Non-coding small nucleolar RNA SNORD17 promotes the progression of hepatocellular carcinoma through a positive feedback loop upon p53 inactivation. Cell Death Differ., 29, 988–1003.

33. Qiu, H., Lee, S., Shang, Y., Wang, W.-Y., Au, K.F., Kamiya, S., Barmada, S.J., Finkbeiner, S., Lui, H., Carlton, C.E., et al. (2014) ALS-associated mutation FUS-R521C causes DNA damage and RNA splicing defects. J. Clin. Invest., 124, 981–999.

34. Reber, S., Stettler, J., Filosa, G., Colombo, M., Jutzi, D., Lenzken, S.C., Schweingruber, C., Bruggmann, R., Bachi, A., Barabino, S.M., et al. (2016) Minor intron splicing is regulated by FUS and affected by ALS-associated FUS mutants. EMBO J., 35, 1504–1521.

35. Orozco, D. and Edbauer, D. (2013) FUS-mediated alternative splicing in the nervous system: consequences for ALS and FTLD. J. Mol. Med. (Berl), 91, 1343–1354.

36. Yamazaki, T., Chen, S., Yu, Y., Yan, B., Haertlein, T.C., Carrasco, M.A., Tapia, J.C., Zhai, B., Das, R., Lalancette-Hebert, M., et al. (2012) FUS-SMN Protein Interactions Link the Motor Neuron Diseases ALS and SMA. Cell Rep., 2, 799–806.

37. Gerbino, V., Carrì, M.T., Cozzolino, M. and Achsel, T. (2013) Mislocalised FUS mutants stall spliceosomal snRNPs in the cytoplasm. Neurobiol. Dis., 55, 120–128.

38. Sun, S., Ling, S.-C., Qiu, J., Albuquerque, C.P., Zhou, Y., Tokunaga, S., Li, H., Qiu, H., Bui, A., Yeo, G.W., et al. (2015) ALS-causative mutations in FUS/TLS confer gain and loss of function by altered association with SMN and U1-snRNP. Nat. Commun., 6, 6171.

39. Kovanda, A., Leonardis, L., Zidar, J., Koritnik, B., Dolenc-Groselj, L., Ristic Kovacic, S., Curk, T. and Rogelj, B. (2018) Differential expression of microRNAs and other small RNAs in muscle tissue of patients with ALS and healthy age-matched controls. Sci. Rep., 8, 5609.

40. Schattner, P., Brooks, A.N. and Lowe, T.M. (2005) The tRNAscan-SE, snoscan and snoGPS web servers for the detection of tRNAs and snoRNAs. Nucleic Acids Res., 33, W686–W689.

41. Bouchard-Bourelle, P., Desjardins-Henri, C., Mathurin-St-Pierre, D., Deschamps-Francoeur, G., Fafard-Couture, É., Garant, J.-M., Elela, S.A. and Scott, M.S. (2020) snoDB: an interactive database of human snoRNA sequences, abundance and interactions. Nucleic Acids Res., 48, D220–D225.

42. Krogh, N., Jansson, M.D., Häfner, S.J., Tehler, D., Birkedal, U., Christensen-Dalsgaard, M., Lund, A.H. and Nielsen, H. (2016) Profiling of 2’-O-Me in human rRNA reveals a subset of fractionally modified positions and provides evidence for ribosome heterogeneity. Nucleic Acids Res., 44, 7884–7895.

43. Diesend, J., Birkedal, U., Kjellin, J., Zhang, J., Jablonski, K.P., Söderbom, F., Nielsen, H. and Hammann, C. (2022) Fractional 2′-O-methylation in the ribosomal RNA of Dictyostelium discoideum supports ribosome heterogeneity in Amoebozoa. Sci. Rep., 12, 1952.

44. Motorin, Y., Quinternet, M., Rhalloussi, W. and Marchand, V. (2021) Constitutive and variable 2’-O-methylation (Nm) in human ribosomal RNA. RNA Biol., 18, 88–97.

45. Khatter, H., Myasnikov, A.G., Natchiar, S.K. and Klaholz, B.P. (2015) Structure of the human 80S ribosome. Nature, 520, 640–645.

46. Lagier-Tourenne, C., Polymenidou, M., Hutt, K.R., Vu, A.Q., Baughn, M., Huelga, S.C., Clutario, K.M., Ling, S.-C., Liang, T.Y., Mazur, C., et al. (2012) Divergent roles of ALS-linked proteins FUS/TLS and TDP-43 intersect in processing long pre-mRNAs. Nat. Neurosci., 15, 1488–1497.

47. Izumikawa, K., Nobe, Y., Ishikawa, H., Yamauchi, Y., Taoka, M., Sato, K., Nakayama, H., Simpson, R.J., Isobe, T. and Takahashi, N. (2019) TDP-43 regulates site-specific 2’-O-methylation of U1 and U2 snRNAs via controlling the Cajal body localization of a subset of C/D scaRNAs. Nucleic Acids Res., 47, 2487–2505.

48. Groen, E.J.N., Fumoto, K., Blokhuis, A.M., Engelen-Lee, J., Zhou, Y., van den Heuvel, D.M.A., Koppers, M., van Diggelen, F., van Heest, J., Demmers, J.A.A., et al. (2013) ALS-associated mutations in FUS disrupt the axonal distribution and function of SMN. Hum. Mol. Genet., 22, 3690–3704.

49. Jutzi, D., Campagne, S., Schmidt, R., Reber, S., Mechtersheimer, J., Gypas, F., Schweingruber, C., Colombo, M., von Schroetter, C., Loughlin, F.E., et al. (2020) Aberrant interaction of FUS with the U1 snRNA provides a molecular mechanism of FUS induced amyotrophic lateral sclerosis. Nat. Commun., 11, 6341.

50. Riva, N., Clarelli, F., Domi, T., Cerri, F., Gallia, F., Trimarco, A., Brambilla, P., Lunetta, C., Lazzerini, A., Lauria, G., et al. (2016) Unraveling gene expression profiles in peripheral motor nerve from amyotrophic lateral sclerosis patients: insights into pathogenesis. Sci. Rep., 6, 39297.

51. Zhao, W., Zhang, S., Zhu, Y., Xi, X., Bao, P., Ma, Z., Kapral, T.H., Chen, S., Zagrovic, B., Yang, Y.T., et al. (2022) POSTAR3: an updated platform for exploring post-transcriptional regulation coordinated by RNA-binding proteins. Nucleic Acids Res., 50, D287–D294.

52. McCann, K.L., Kavari, S.L., Burkholder, A.B., Phillips, B.T. and Hall, T.M.T. (2020) H/ACA snoRNA levels are regulated during stem cell differentiation. Nucleic Acids Res., 48, 8686–8703.

53. Balogh, E., Chandler, J.C., Varga, M., Tahoun, M., Menyhárd, D.K., Schay, G., Goncalves, T., Hamar, R., Légrádi, R., Szekeres, Á., et al. (2020) Pseudouridylation defect due to DKC1 and NOP10 mutations causes nephrotic syndrome with cataracts, hearing impairment, and enterocolitis. Proc. Natl. Acad. Sci. U. S. A., 117, 15137–15147.

54. Kamelgarn, M., Chen, J., Kuang, L., Jin, H., Kasarskis, E.J. and Zhu, H. (2018) ALS mutations of FUS suppress protein translation and disrupt the regulation of nonsense-mediated decay. Proc. Natl. Acad. Sci. U. S. A., 115, E11904–E11913.

55. Sahoo, T., del Gaudio, D., German, J.R., Shinawi, M., Peters, S.U., Person, R.E., Garnica, A., Cheung, S.W. and Beaudet, A.L. (2008) Prader-Willi phenotype caused by paternal deficiency for the HBII-85 C/D box small nucleolar RNA cluster. Nat. Genet., 40, 719–721.

56. Duker, A.L., Ballif, B.C., Bawle, E.V., Person, R.E., Mahadevan, S., Alliman, S., Thompson, R., Traylor, R., Bejjani, B.A., Shaffer, L.G., et al. (2010) Paternally inherited microdeletion at 15q11.2 confirms a significant role for the SNORD116 C/D box snoRNA cluster in Prader–Willi syndrome. Eur. J. Hum. Genet., 18, 1196–1201.

57. Valleron, W., Laprevotte, E., Gautier, E.-F., Quelen, C., Demur, C., Delabesse, E., Agirre, X., Prósper, F., Kiss, T. and Brousset, P. (2012) Specific small nucleolar RNA expression profiles in acute leukemia. Leukemia, 26, 2052–2060.

58. Reber, S., Mechtersheimer, J., Nasif, S., Benitez, J.A., Colombo, M., Domanski, M., Jutzi, D., Hedlund, E. and Ruepp, M.-D. (2018) CRISPR-Trap: a clean approach for the generation of gene knockouts and gene replacements in human cells. Mol. Biol. Cell, 29, 75–83.

59. Ran, F.A., Hsu, P.D., Wright, J., Agarwala, V., Scott, D.A. and Zhang, F. (2013) Genome engineering using the CRISPR-Cas9 system. Nat. Protoc., 8, 2281–2308.

60. Simpson, P.B., Bacha, J.I., Palfreyman, E.L., Woollacott, A.J., McKernan, R.M. and Kerby, J. (2001) Retinoic acid evoked-differentiation of neuroblastoma cells predominates over growth factor stimulation: an automated image capture and quantitation approach to neuritogenesis. Anal. Biochem., 298, 163–169.

61. LaMar, D. (2015) FastQC..

62. Gordon, A. (2022) agordon/fastx_toolkit. agordon/fastx_toolkit.

63. Langmead, B. and Salzberg, S.L. (2012) Fast gapped-read alignment with Bowtie 2. Nat. Methods, 9, 357–359.

64. Li, H., Handsaker, B., Wysoker, A., Fennell, T., Ruan, J., Homer, N., Marth, G., Abecasis, G., Durbin, R., and 1000 Genome Project Data Processing Subgroup (2009) The Sequence Alignment/Map format and SAMtools. Bioinformatics, 25, 2078–2079.

65. Liao, Y., Smyth, G.K. and Shi, W. (2014) featureCounts: an efficient general purpose program for assigning sequence reads to genomic features. Bioinformatics, 30, 923–930.

66. Love, M.I., Huber, W. and Anders, S. (2014) Moderated estimation of fold change and dispersion for RNA-seq data with DESeq2. Genome Biol., 15, 550.

67. gplots package - RDocumentation https://rdocumentation.org/packages/gplots/versions/3.1.1 (accessed Mar 30, 2022).

68. Blighe, K. (2022) EnhancedVolcano: publication-ready volcano plots with enhanced colouring and labeling.

