## Supplemental Data for "FUS modulates the level of ribosomal RNA modifications by regulating a subset of snoRNA expression"

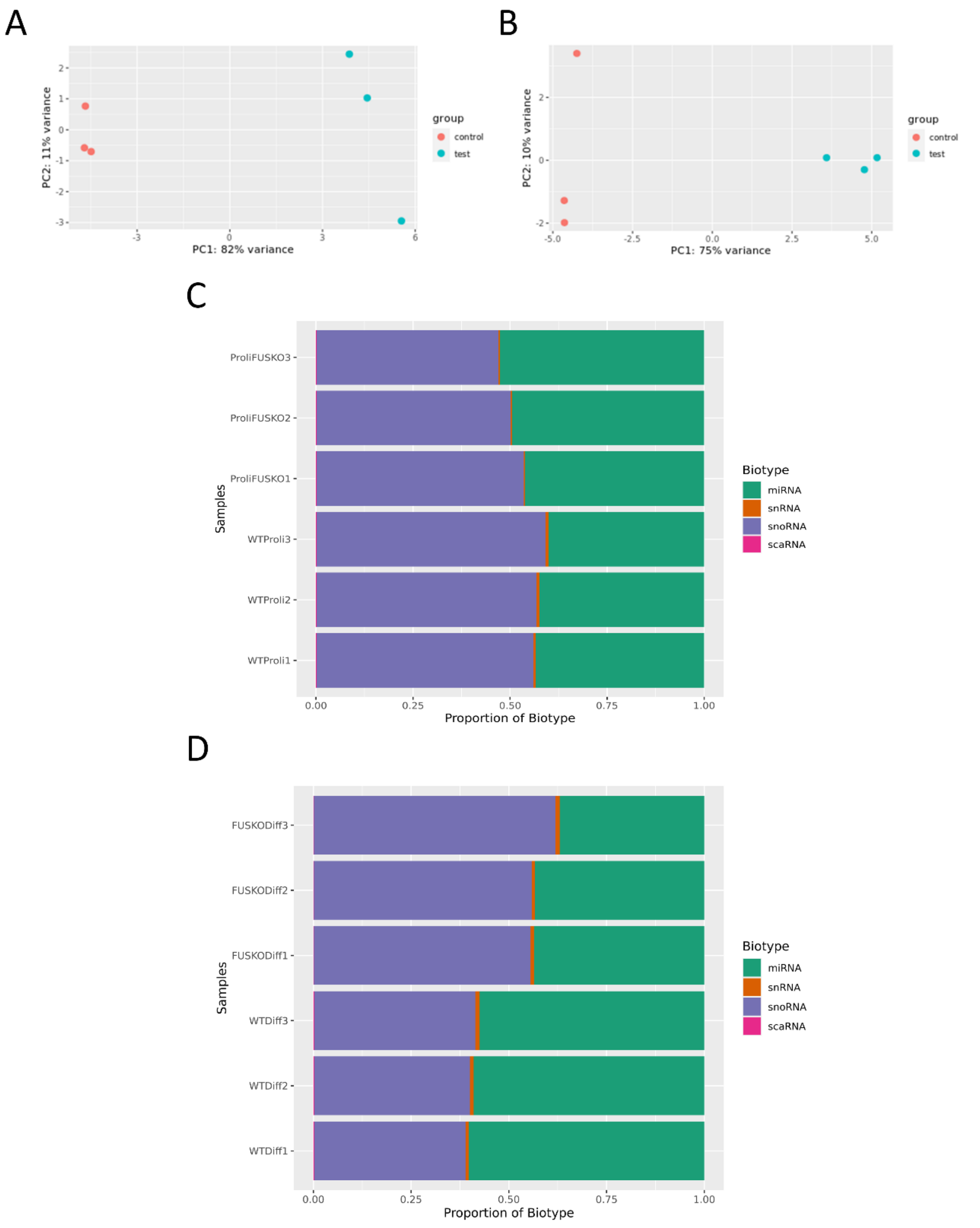


**Supplementary Fig. S1**. PCA (principal component analysis) plots generated using ggplot2 library in R. Raw count matrix generated from FeatureCounts was used with rlog (regularized log) transformation in DESeq2 R package. Control=wild-type cells, test=FUS KO cells (A and B). Proportion of small RNA biotypes counted by the FeatureCounts tool and plotted using libraries ggplot2 and reshape2 in R. X-axis denotes proportion of each biotype in the samples and on Y-axis are the sample names with three biological replicates each (C and D).

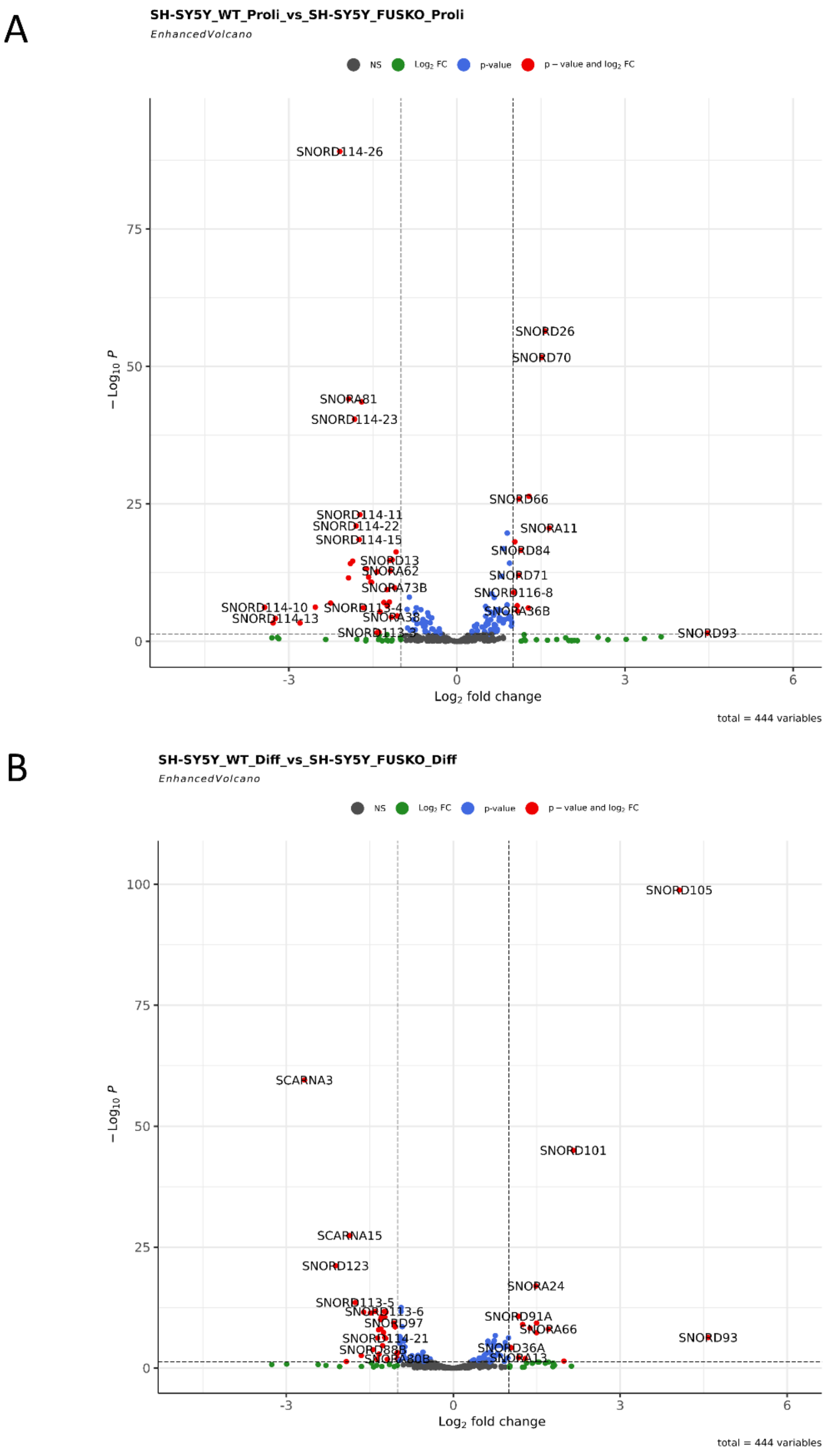


**Supplementary Fig. S2**. Volcano plots for proliferating (A) and differentiated (B) SH-SY5Y cells. Enhanced Volcano plots were generated using Enhanced Volcano R package. Raw count matrix with only snoRNAs was used in DESeq2 to get differentially expressed snoRNAs (padj < 0.05), the tabular file generated was then used in Enhanced Volcano library with x-axis = 'log2FoldChange' and y-axis = 'padj'. Representative differentially expressed snoRNAs are annotated. (NS: non-significant).

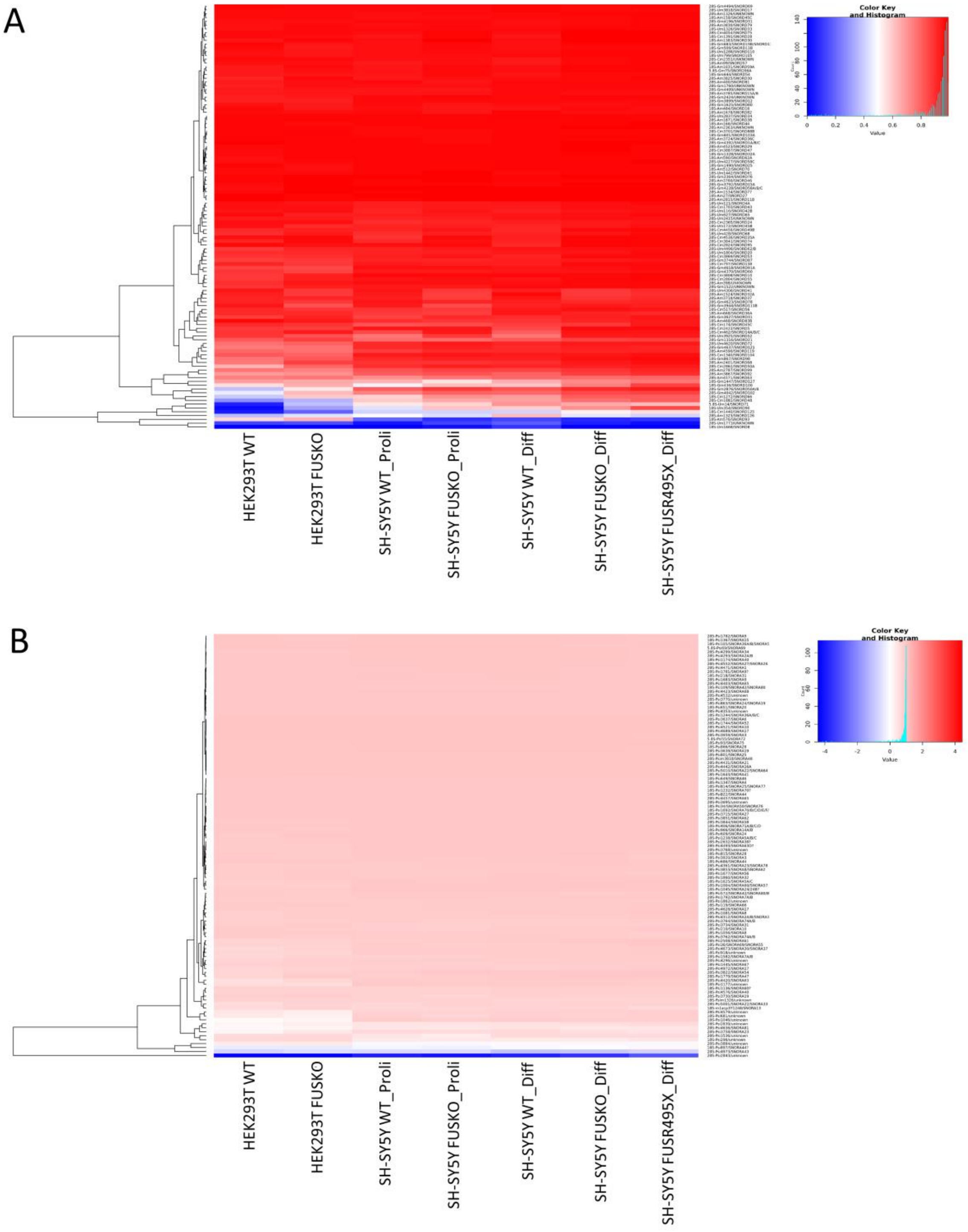


**Supplementary Fig. S3.** Heatmaps representing the average 2’-O-Me (A) and pseudouridine (B) levels in all cells analyzed (HEK293T WT and FUS KO, SH-SY5Y proliferating (Proli) WT and FUS KO, SH-SY5Y differentiated (Diff) WT, FUS KO and FUS R495X), representing the average of 3 biological replicates (N=3) from each cell type. The blue and red colors show a low and high proportion of modification, respectively.

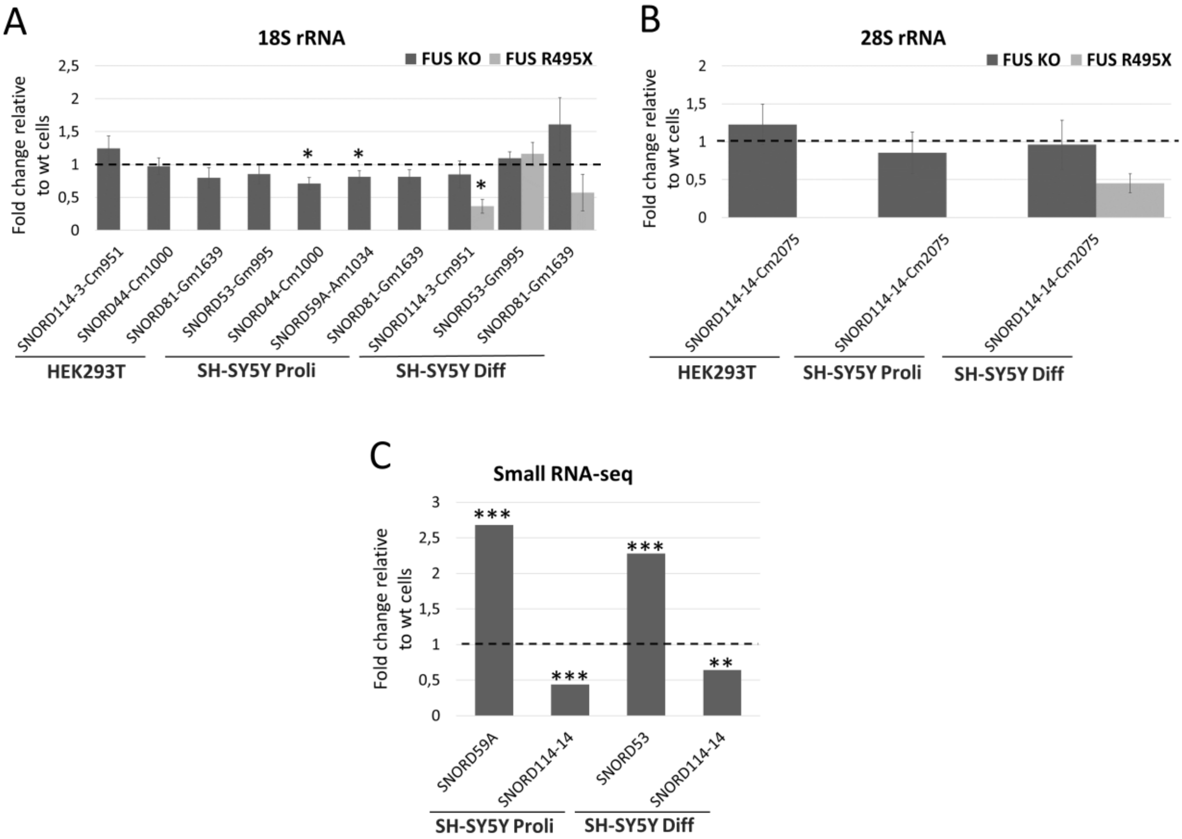


**Supplementary Fig. S4.** RT-qPCR analysis of relative expression level of selected C/D box snoRNAs predicted to guide methylation at putative new positions in 18S rRNA (A) and 28S rRNA (B). Small RNA-seq results of relative expression levels of selected snoRNAs that guide methylation and pseudouridylation of residues in 18S rRNA and 28S rRNA in SH-SY5Y proliferating and differentiated cells (C). HEK293T FUS KO cells, proliferating (Proli), differentiated (Diff) SH-SY5Y FUS KO cells, and FUS R495X cells were compared to WT cells, respectively. Error bars represent the SD of three biological replicates (N=3). P-values were calculated using Student's t-test, and the statistical significance is defined as follows: $*P \leq 0.05$; $**P \leq 0.01$; $***P \leq 0.001$.

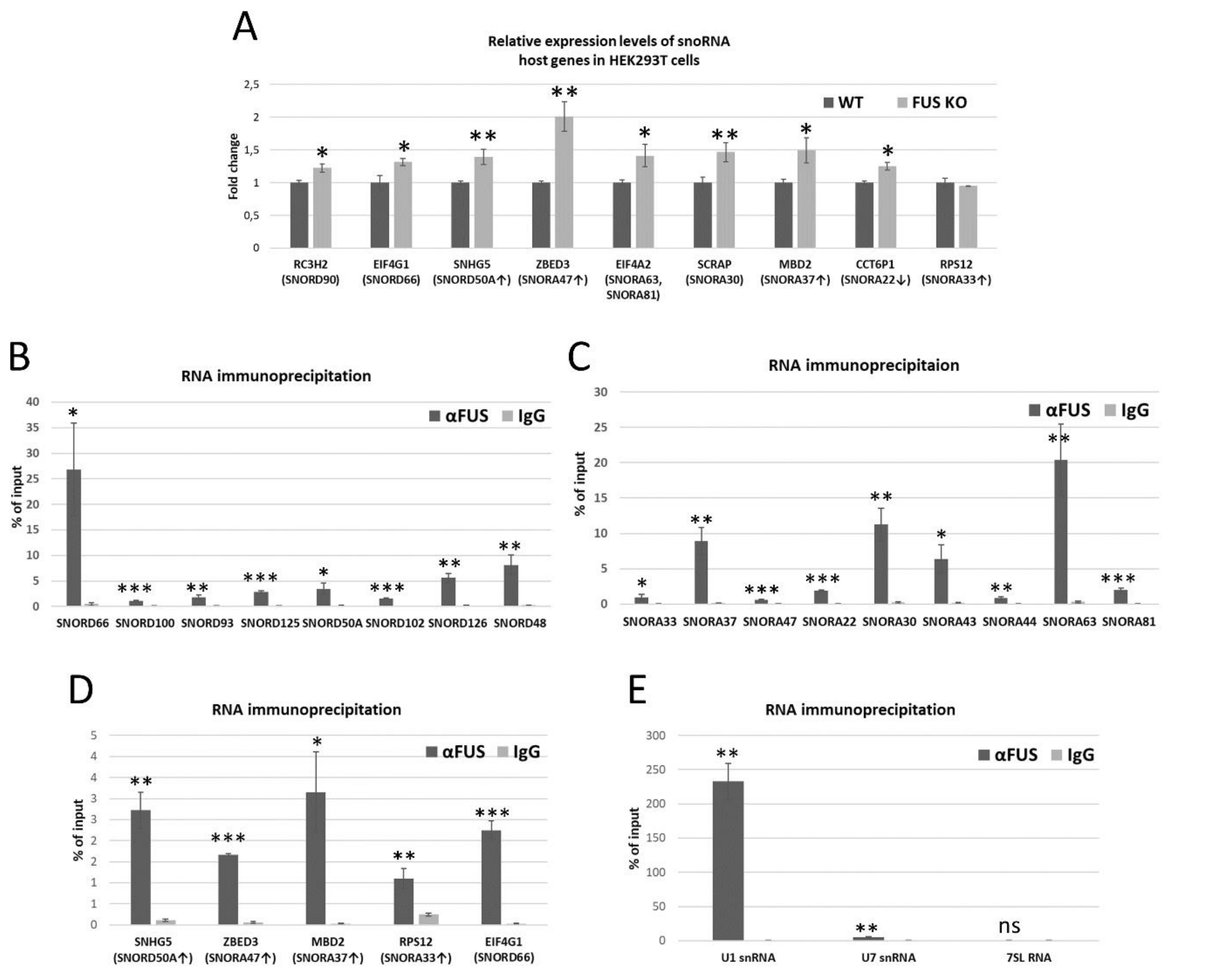


**Supplementary Fig. S5.** RT-qPCR analysis of relative expression levels of host genes of selected snoRNAs in HEK293T WT and FUS KO cells (A). RNA immunoprecipitation (RIP) experiment to test whether FUS can bind C/D box snoRNAs (B), H/ACA box snoRNAs (C), and snoRNA-host gene transcripts (D) in HEK293T cells. 7SL RNA was used as a negative control, U1 snRNA and U7 snRNAs were used as a positive control (E). Error bars represent the SD of three biological replicates (N=3). P-values were calculated using Student's t-test, and the statistical significance is defined as follows: *P ≤ 0.05; **P ≤ 0.01; ***P ≤ 0.001.

RT-qPCR

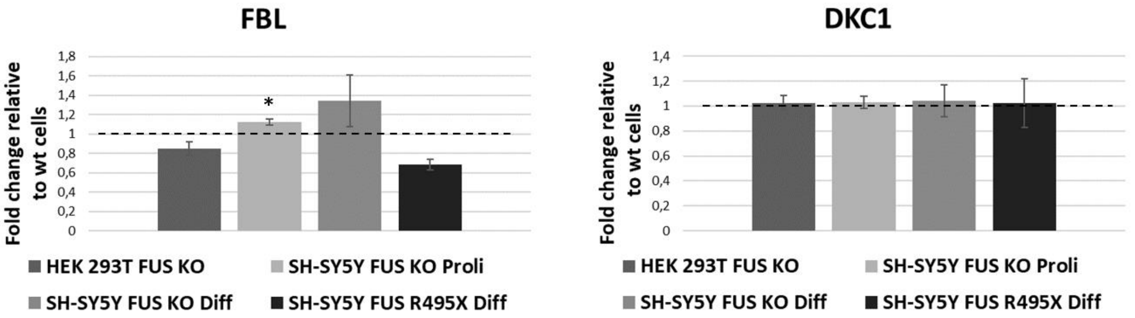


Western blot

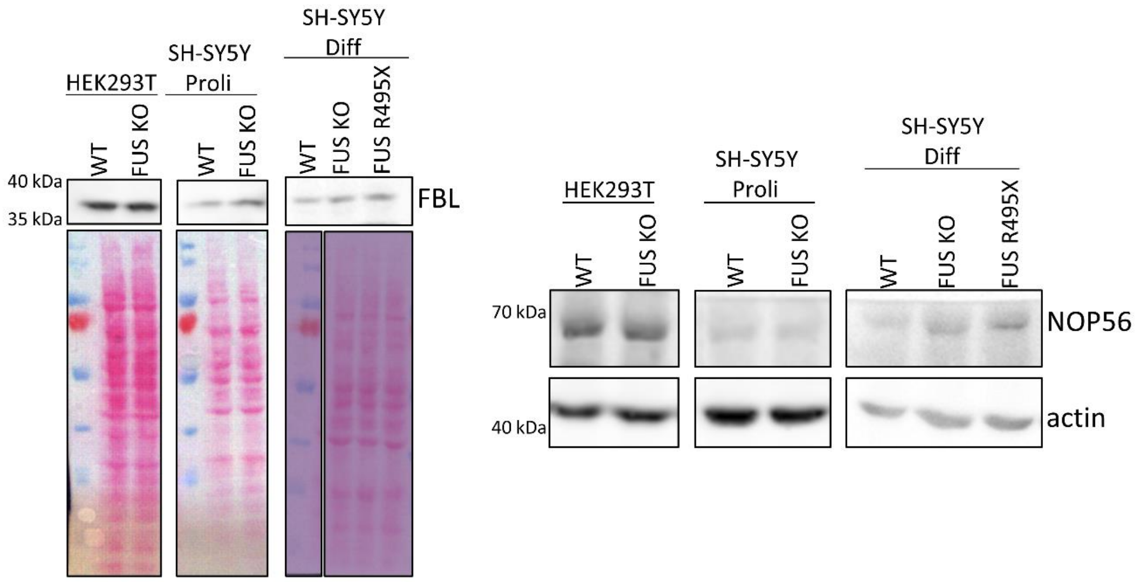


**Supplementary Fig. S6.** RT-qPCR and Western blot and immunodetection to analyze the expression of fibrillarin (FBL), NOP56, and dyskerin (DKC1) in FUS KO cells and FUS R495X cells in comparison to WT cells. Error bars represent the SD of three biological replicates (N=3). P-values were calculated using Student's t-test, and the statistical significance is defined as follows: $^*P \leq 0.05$. For staining, anti-FIB, anti-NOP56 and anti-actin antibodies were used. Actin level or Ponceau S-stained membranes were used as loading controls. All experiments were done in 3 biological replicates.

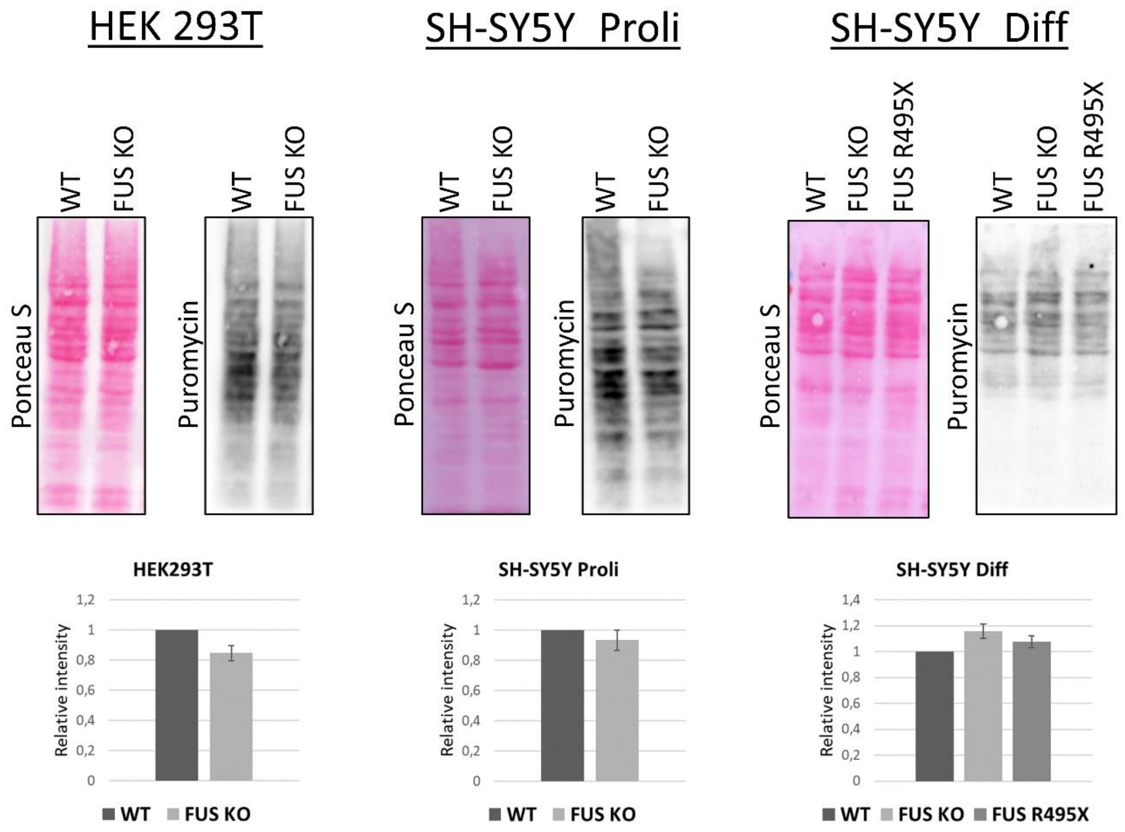


**Supplementary Fig. S7.** SUnSET assay to analyze global translation efficiency in all cells analyzed (HEK293T WT and FUS KO, SH-SY5Y proliferating (Proli) WT and FUS KO, SH-SY5Y differentiated (Diff) WT, FUS KO and FUS R495X). Ponceau S-stained membranes were used as loading controls. The graphs represent the average of three biological replicates (N=3) and were performed using the intensity of puromycinylated proteins normalized to Ponceau S signal in each lane. Error bars represent the SD of three biological replicates.

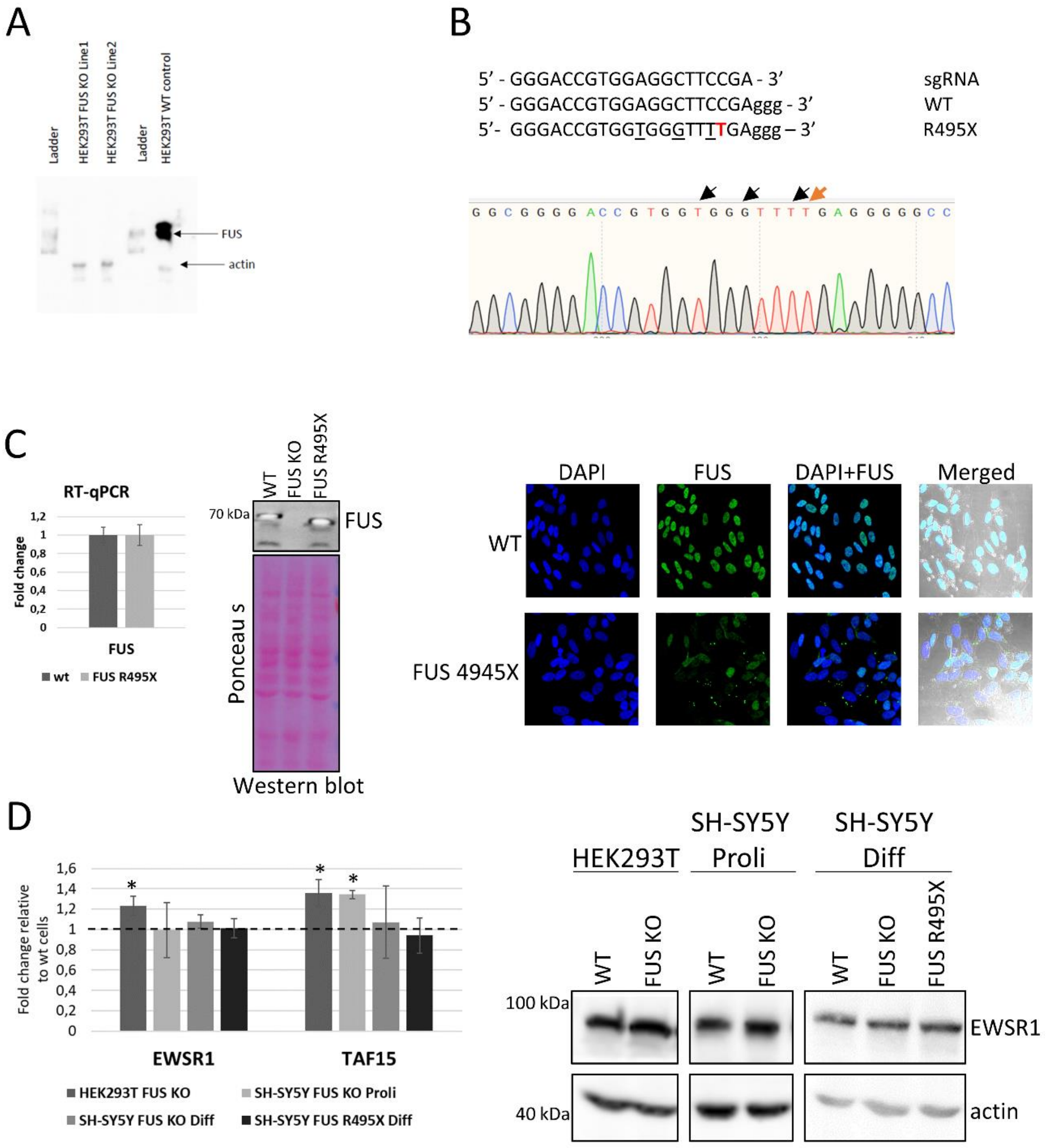


**Supplementary Fig. S8.** Western blot and immunodetection to confirm the knockout of *FUS* gene in HEK293T cells. For staining, anti-FUS and anti-actin antibodies were used with dilutions as mentioned in the methodology section. HEK293T FUS KO cells show no band corresponding to FUS protein (A). Representative sequence alignment and sequencing chromatogram of CRISPR/Cas9-induced FUS R495X mutation (arginine to stop codon, marked in red). Small letters correspond to PAM sequence. Silent mutations in sgRNA seeding regions are underlined (B). RT-qPCR, Western blot and immunodetection as well as immunofluorescence to confirm the expression and partial cytoplasmic mislocalization of FUS R495X in differentiated SH-SY5Y cells (C). RT-qPCR and Western blot and immunodetection to test the expression level of TAF15 and EWSR1 in all cells analyzed (HEK293T WT and FUS KO, SH-SY5Y proliferating (Proli) WT and FUS KO, SH-SY5Y differentiated (Diff) WT, FUS KO and FUS R495X) (D). Error bars represent the SD of three biological replicates (N=3). P-values were calculated using Student's t-test, and the statistical significance is defined as follows: $^*P \leq 0.05$. All experiments were done in three biological replicates.

**Supplementary Table S3.** Representative results from Snoscan.

snoDB database fasta file was provided to the Snoscan tool with human ribosomal RNAs SSU and LSU as target RNAs. Snoscan was able to identify known 2'-O-Me sites, one example is mentioned here (28S-Cm2421). Alignment of putative new 2'-O-Me sites with guide snoRNAs predicted by the Snoscan tool are mentioned below:

**1. 18S-Cm951 – Putative guide C/D box snoRNA SNORD114-3/12/14**

```
>> SNORD114-3  13.25  (10-76)  Cmpl: NU_167214.1:109078-110946-Cm951 (-)  12/1 bp  Gs-DpBox: 20 (11)  Len: 67  TS

No known meth site found      Guide Seq Sc: -4.33  (15.04 -2.68 -15.69 -1.00)
                                 *
Db seq:  5'-                  CGCCGCUAGAGGU -3'  NU_167214.1:109078-110946  (948-956)
                              ||||:|:|| |||
Qry seq: 3'-             AGUUUGCGGUGGUCACCA -5'  SNORD114-3  (32-20)

Strong terminal stem:           +-[C Box] -N- CCAGGU-D - 5'                Stem Sc: 5.76 (6 bp)
                         |             ||||||
                         +---[D Box] - GGUCCAA - 3'          Stem Transit Sc: -0.63
>Summary      [ C Box ] --         -- [ Cmpl/ Mism ]  X [D'Bx] --       -- [D Bx]  Length
>Meth Cm 951  [AUGAUGA] --   0 bp  -- [  12 / 1    ]  1 [UUGA] -- 33 bp -- [CUGA]   67 bp
>Sc    13.25  [ 11.42 ] --  -6.55  -- [ 15.04 bits ]    [3.67]    -3.54    [8.05]
Candidate sequence:
>SNORD114-3  13.25  (10-76)  Cmpl: NU_167214.1:109078-110946-Cm951  Len: 67
ACCAATGATGACCACTGGTGGCGTTTGAGTCATGGACGATGAATACTACGTGTCTGAAACTCTGAGG
```

**2. 18S-Gm995 – Putative guide C/D box snoRNA SNORD53**

```
>> SNORD53  15.45  (7-81)  Cmpl: NU_167214.1:109078-110946-Gm995 (-)  12/2 bp  Gs-DpBox: 22 (16)  Len: 75  TS
No known meth site found      Guide Seq Sc: -1.19  (18.18 -2.68 -15.69 -1.00)
                                *
Db seq:  5'-                 AGCGAAAGCAUUUG -3'  NU_167214.1:109078-110946  (992-1005)
                             ||||||| ||| ||
Qry seq: 3'-            GGUCGUCGCUUUGGUAUAC -5'   SNORD53    (35-22)

Terminal stem:                  +-[C Box] -N- CGUADRON - 5'                Stem Sc: 2.31 (4 bp)
                         |             ||||
                         +---[D Box] - GCAU - 3'             Stem Transit Sc: -0.63
```

```
>Summary      [ C Box ] --         -- [ Cmpl/ Mism ]  X [D'Bx] --       -- [D Bx]  Length
>Meth Gm 995  [AUGAUGA] --   4 bp  -- [  12 / 2    ]  1 [CUGG] -- 35 bp -- [CUGA]   75 bp
>Sc    15.45  [ 11.42 ] --  -3.32  -- [ 18.18 bits ]    [2.94]    -3.54    [8.05]
Candidate sequence:
>SNORD53  15.45  (7-81)  Cmpl: NU_167214.1:109078-110946-Gm995  Len: 75
TGCTATGATGACATCCATATGGTTTCGCTGCTGGCTGAGTTTCAGAGATGACACCTTTCTCTTGGCTGTCTGAGC
```

**3. 18S-Cm1000 - Putative guide C/D box snoRNA SNORD44**

```
>> SNORD44  12.92  (2-61)  Cmpl: Hu-18S-Cm1000 (-)  10/2 bp  Gs-DpBox: 16 (15)  Len: 60

No known meth site found      Guide Seq Sc: -5.42  (14.60 -1.98 -15.69 -2.35)

                                    *
Db seq:  5'-                   GAAAGCAUUUGC -3'    Hu-18S    (995-1006)
                               |  |||||||||
Qry seq: 3'-                AGUCAGUCGUAAACG -5'   SNORD44    (27-16)

No terminal stem:              +-[C Box] -N- GUCC - 5'             Stem Sc: -1.71 (1 bp)
                         |             |
                         +---[D Box] - CU - 3'               Stem Transit Sc: -1.50

>Summary      [ C Box ] --         -- [ Cmpl/ Mism ]  X [D'Bx] --       -- [D Bx]  Length
>Meth Cm1000  [AUGAUGA] --   3 bp  -- [  10 / 2    ] -1 [CUGA] -- 25 bp -- [CUGA]   60 bp
>Sc    12.92  [ 11.42 ] --  -3.32  -- [ 14.60 bits ]    [7.09]    -2.80    [8.05]

Candidate sequence:
>SNORD44  12.92  (2-61)  Cmpl: Hu-18S-Cm1000  Len: 60
CTGGATGATGATAAGCAAATGCTGACTGAACATGAAGGTCTTAATTAGCTCTAACTGACT
```

**4. 18S-Am1034 - Putative guide C/D box snoRNA SNORD59A**

```
>> SNORD59A  12.91  (3-72)  Cmpl: Hu-18S-Am1034 (-)  10/0 bp  Gs-DpBox: 26 (24)  Len: 70  TS

No known meth site found      Guide Seq Sc: -0.55  (18.91 -2.06 -15.69 -1.72)

                                     *
Db seq:  5'-                     AACGAAAGUC -3'    Hu-18S    (1030-1042)
                                 ||||||||||
Qry seq: 3'-                 CUUCUUGCUUUCAG -5'  SNORD59A    (35-26)
```

```
Terminal stem:                  +-[C Box] -N- CUUCC - 5'            Stem Sc: 4.29 (5 bp)
                         |             |||||
                         +---[D Box] - GAAGG - 3'            Stem Transit Sc: -0.63

>Summary      [ C Box ] --         -- [ Cmpl/ Mism ]  X [D'Bx] --       -- [D Bx]  Length
>Meth Am1034  [AUGAUGA] --  12 bp  -- [  10 / 0    ]  0 [CUUC] -- 27 bp -- [CUGA]   70 bp
>Sc    12.91  [ 11.42 ] --  -3.97  -- [ 18.91 bits ]    [-1.57]    -3.54    [8.05]

Candidate sequence:
>SNORD59A  12.91  (3-72)  Cmpl: Hu-18S-Am1034  Len: 70
TTCTATGATGATTTTATCAAAATGACTTTCGTTCTTCTGAGTTTGCTGAAGCCACATTTA
GGTACTGAGA
```

**4. 18S-Am1639 - Putative guide C/D box snoRNA SNORD81**

```
>> SNORD81  12.30  (5-72)  Cmpl: Hu-18S-Gm1639 (-)  9/1 bp  Gs-DpBox: 30 (26)  Len: 68  TS

No known meth site found      Guide Seq Sc: -7.33  (11.42 -2.06 -15.69 -1.00)

                                   *
Db seq:  5'-                    GAGGGAAUUC -3'    Hu-18S    (1636-1646)
                                |||:|| |||
Qry seq: 3'-               AGUCACUCUCUCAAG -5'   SNORD81    (39-30)

Strong terminal stem:           +-[C Box] -N- AUAAGAC - 5'          Stem Sc: 6.14 (7 bp)
                         |             |||||||
                         +---[D Box] - UAUUCUG - 3'          Stem Transit Sc: -0.63

>Summary      [ C Box ] --         -- [ Cmpl/ Mism ]  X [D'Bx] --       -- [D Bx]  Length
>Meth Gm1639  [AUGAUGA] --  14 bp  -- [   9 / 1    ]  1 [CUGA] -- 22 bp -- [CUGA]   68 bp
>Sc    12.30  [ 11.42 ] --  -8.14  -- [ 11.42 bits ]    [7.09]    -3.69    [8.05]

Candidate sequence:
>SNORD81  12.30  (5-72)  Cmpl: Hu-18S-Gm1639  Len: 68
ATACATGATGATCTCAATCCAACTTGAACTCTCTCACTGATTACTTGATGACAATAAAAT
ATCTGATA
```

**5. 28S-Cm2075 - Putative guide C/D box snoRNA SNORD114-14**

```
>> SNORD114-14  22.70  (10-76)  Cmpl: NU_167214.1:113348-118417-Cm2084 (-)  10/0 bp  Gs-DpBox: 23 (14)  Len: 67  TS
No known meth site found      Guide Seq Sc: 2.74  (21.49 -2.06 -15.69 -1.00)
```

```
                                   *
Db seq:  5'-                    CGCCGGCAGU -3'  NU_167214.1:113348-118417  (2081-2089)
                                ||||||||||
Qry seq: 3'-               AGUAUGCGGCCGUCA -5'  SNORD114-14 (32-23)

Strong terminal stem:           +-[C Box] -N- CCAGGU-D - 5'                Stem Sc: 5.76 (6 bp)
                         |             ||||||
                         +---[D Box] - GGUCCA - 3'           Stem Transit Sc: -0.63
>Summary      [ C Box ] --         -- [ Cmpl/ Mism ]  X [D'Bx] --       -- [D Bx]  Length
>Meth Cm2084  [AUGAUGA] --   2 bp  -- [  10 / 0    ]  1 [AUGA] -- 33 bp -- [CUGA]   67 bp
>Sc    22.70  [ 11.42 ] --  -3.32  -- [ 21.49 bits ]    [2.81]    -3.54    [8.05]
Candidate sequence:
>SNORD114-14  22.70  (10-76)  Cmpl: NU_167214.1:113348-118417-Cm2084  Len: 67
ACCAATGATGACAACTGCCGGCGTATGAGTGTTGGGTGATGAATAATACGTGTCTAGAACTCTGAGG
```

**4. 28S-Cm2421 – Guide C/D box snoRNA SNORD5**

```
>> SNORD5  19.01  (8-75)  Cmpl: NU_167214.1:113348-118417-Cm2422 (-)  13/0 bp  Gs-DpBox: 26 (19)  Len: 68

No known meth site found      Guide Seq Sc: 5.68  (25.05 -2.68 -15.69 -1.00)
                                *
Db seq:  5'-                 CAGCAGUUGAACA -3'  NU_167214.1:113348-118417  (2419-2433)
                             |||||||||||||
Qry seq: 3'-            AGUAAGUCGUCAACUUGU -5'    SNORD5    (38-26)

Possible terminal stem:         +-[C Box] -N-  ACUUGDRO - 5'               Stem Sc: -0.19 (3 bp)
                         |              ||:
                         +---[D Box] - CAGAG - 3'            Stem Transit Sc: -1.50
>Summary      [ C Box ] --         -- [ Cmpl/ Mism ]  X [D'Bx] --       -- [D Bx]  Length
>Meth Cm2422  [AUGAUGA] --   7 bp  -- [  13 / 0    ]  1 [AUGA] -- 26 bp -- [CUGA]   68 bp
>Sc    19.01  [ 11.42 ] --  -3.32  -- [ 25.05 bits ]    [2.81]    -3.54    [8.05]
Candidate sequence:
>SNORD5  19.01  (8-75)  Cmpl: NU_167214.1:113348-118417-Cm2422  Len: 68
TCAGATGATGAATTTAACTGTTCAACTGCTGAATGATAACGGGCATGAACTAAAACTTAATTCTGAC
```

**Supplementary Table S5.** Primers used in PCR and RT-qPCR

| | |
|---|---|
| AGCTGCGGTTTCAGGTAGTC | FUS (Fwd) |
| ACTTTTAATGGGAACCAGAGGT | FUS (Rev) |
| GTCTAATGATGAATTTCATAGGGCA | SNORD90 (Fwd) |
| GTCTTCAGATTCCACAGTAGGAG | SNORD90 (Rev) |
| CTGATGACTTCCTGTTAGTGCC | SNORD66 (Fwd) |
| TCCTCAGATCCTCAGTTCCATC | SNORD66 (Rev) |
| ACATGATGACAACTGGCTCCC | SNORD100 (Fwd) |
| GCTGTAATCAGAAGGGTGACAT | SNORD100 (Rev) |
| TGGCCAAGGATGAGAACTCTA | SNORD93 (Fwd) |
| GGCCTCAGGTAAATCCTTTAATCC | SNORD93 (Rev) |
| AGCCCCTCCTGATGATTC | SNORD125 (Fwd) |
| TTCAGTCAACTTCTTAGAGGCTC | SNORD125 (Rev) |
| TGTGATGATCTTATCCCGAACCT | SNORD50A (Fwd) |
| ATCTCAGAAGCCAGATCCGT | SNORD50A (Rev) |
| AGCTTAATGATGACTGTTTTTTTTGATTGCTTGA | SNORD102 (Fwd) |
| AGCTTTCAGAGCCGGTGAAATGTGTTTTTC | SNORD102 (Rev) |
| CATGATGAAATGCATGTTAAGTCCGT | SNORD126 (Fwd) |
| GCTCAGAGCATGTGTTTAATCAGGC | SNORD126 (Rev) |
| GATGATGACCCCAGGTAACTCTTG | SNORD48 (Fwd) |
| GTCAGAGCGCTGCGGTGAT | SNORD48 (Rev) |
| TGGTGCTGTGATGATGCCTTA | SNORD92 (Fwd) |
| GCTCAGACACAGCCAAGGAA | SNORD92 (Rev) |
| CAATGATGACCACTGGTGGCG | SNORD114-3 (Fwd) |
| TTGGACCTCAGAGTTTCAGACA | SNORD114-3 (Rev) |
| CCTGGATGATGATAAGCAAATGCTGACT | SNORD44 (Fwd) |
| AGTCAGTTAGAGCTAATTAAGACCTTCATG | SNORD44 (Rev) |
| TGATGACATCCATATGGTTTCGCTG | SNORD53 (Fwd) |
| GCTCAGACAGCCAAGAGAAAG | SNORD53 (Rev) |
| CCTTCTATGATGATTTTATCAAAATGACTTTCGTT | SNORD59A (Fwd) |
| CCTTCTCAGTACCTAAATGTGGCTTCA | SNORD59A (Rev) |
| CAGAATACATGATGATCTCAATCCAACTTGAAC | SNORD81 (Fwd) |
| CAGAATATCAGATATTTTATTGTCATCAAGTAATCAGTG | SNORD81 (Rev) |
| GACCAATGATGACAACTGCC | SNORD114-14 (Fwd) |
| GACCTCAGAGTTCTAGACACGTATT | SNORD114-14 (Rev) |
| GTGCTGTGTTGTCGTTCCCC | SNORA63 (Fwd) |
| GCTGCTACAGGAGAATAGCAGA | SNORA63 (Rev) |
| CACTTTCACAGTTCCTTCCCC | SNORA30 (Fwd) |
| TCAAGGGTTTTCTCTCAGCACC | SNORA30 (Rev) |
| AGCACTTTCACAGGTCCTCCC | SNORA37 (Fwd) |
| GGCAAGGATGCCAACAAAGGT | SNORA37 (Rev) |
| TTGCACAGTGAACACCCAAGT | SNORA22 (Fwd) |
| CAGAGGAGAAGAGCAGGCAAT | SNORA22 (Rev) |
| AGCCAGCCAATGAATCTGCTT | SNORA33 (Fwd) |
| AGGCTCGTAACATGGCTTTACT | SNORA33 (Rev) |
| CCTTCCACCGGTTAAGACCTC | SNORA47 (Fwd) |

| | |
|---|---|
| CAAATGTCGGCCAGCACAGC | SNORA47 (Rev) |
| TTCGTAACCCGTTAGCCTGG | SNORA54 (Fwd) |
| AGTCAGTCATGTGTCGCTGG | SNORA54 (Rev) |
| GCTTCGGAAAGGGAGGGAAA | SNORA7A/7B (Fwd) |
| CTGTCGCAGAGTGTCTTCCA | SNORA7A/7B (Rev) |
| CTCCAACTGCATGCAAGAGC | SNORA44 (Fwd) |
| ATAGGAAAGCTGAGTGGCAGC | SNORA44 (Rev) |
| ATTGCAGACACTAGGACCATGT | SNORA81 (Fwd) |
| AGAAAGAGGTCCACCCCAGT | SNORA81 (Rev) |
| GTTGGCACCACAGACAGTTG | SNORA43 (Fwd) |
| AAACCATTCTCAGTGCCCAC | SNORA43 (Rev) |
| GGTGGAGGAAGAAGGTCGTG | RC3H2 (For) |
| CTGACAGCGGCCCATAGATT | RC3H2 (Rev) |
| ACCTGAGGAACTGCTCAACG | EIF4G1 (For) |
| AAGGAGCCGTAGCTGGAGTA | EIF4G1 (Rev) |
| GCTGGAGGTGTAATGGACG | RPS12 (For) |
| TTCGCGAATTCCACGTGCT | RPS12 (Rev) |
| CTTAATTGGGGCGGAGGGTT | SNHG5 (For) |
| ATCCGAATTGCACACAACGC | SNHG5 (Rev) |
| AGAGCGGGAAGAGGATGGAT | MBD2 (For) |
| TCGCTCTTGCCAGCACTTAG | MBD2 (Rev) |
| GAATGCAGCGCTGTGTCTTT | ZBED3 (For) |
| CCGAGATGGTAGATCCCCCT | ZBED3 (Rev) |
| GCTGGCGTTCAACATTAGCG | CCT6P1 (For) |
| ATCTTTATGGTCCCTTTGGGC | CCT6P1 (Rev) |
| GCTCTGACTGTGAACCAGAGG | SRCAP (For) |
| TGACTGACTGCTACTATCCTCC | SRCAP (Rev) |
| AGCAACTGGAATGAGATTGTTGA | EIF4A2 (For) |
| CTGCTGAATAGCGGAAGGCT | EIF4A2 (Rev) |
| CTGTAGTGCGCTATGCCGAT | 7SL RNA (For) |
| CACGGGAGTTTTGACCTGCT | 7SL RNA (Rev) |
| ACTCCAGTTATGGACAAAGTCAGT | TAF15 (For) |
| TGGCTGGTCATAGGAAGGTG | TAF15 (Rev) |
| ATGGCGTCCACGGATTACAG | EWSR1 (For) |
| CCATATGCCTGGGTGGTCTG | EWSR1 (Rev) |
| GGCGGATGCGGAAGTAAT | DKC1 (For) |
| CCACTGAGACGTGTCCAACTT | DKC1 (Rev) |
| GAGGCTTCCATTCTGGTGGCAA | FBL (For) |
| CAGGTTCTTGGTGACCAGTGCA | FBL (Rev) |
| CAGTGTTACAGCTCTTTTAGAATTTG | U7 snRNA (For) |
| TTCCGGTAAAAAGCCAGAAA | U7 snRNA (Rev) |
| GATACCATGATCACGAAGGTGGTT | U1 snRNA (For) |
| CACAAATTATGCAGTCGAGTTTCC | U1 snRNA (Rev) |
| CTCAACGACCACTTTGTCAAGCT | GAPDH (For) |
| TCTTACTCCTTGGAGGCCATGT | GAPDH (Rev) |
